# Sapphire-Supported Nanopores for Low-Noise DNA Sensing

**DOI:** 10.1101/2020.03.02.973826

**Authors:** Pengkun Xia, Jiawei Zuo, Pravin Paudel, Shinhyuk Choi, Xiahui Chen, Weisi Song, JongOne Im, Chao Wang

## Abstract

Silicon-supported (SiS) solid-state nanopores have broad applications in single-molecule biosensing and diagnostics, but their high capacitive noise has seriously limited both their sensing accuracy and recording speed. Nanopores on insulating glass have demonstrated reduced capacitance and noise, but it remains challenging to bulk-etch amorphous glass to create membranes reproducibly and uniformly. Here a new approach is reported to form triangular sapphire-supported (SaS) nanopore membranes by batch-processing-compatible anisotropic wet etching of sapphire, with membrane dimension demonstrated from ~200 μm to 5 μm. A SaS nanopore in 68 μm-wide silicon nitride membrane has 130 times smaller capacitance (10 pF) compared to a SiS nanopore (~4 μm SiN membrane, ~1.3 nF), despite a 100 times larger membrane. It has a current noise of 18 pA over 100 kHz bandwidth, much smaller than that from our SiS nanopore (46 pA) and comparable with the best reported low-noise nanopores. Further, the SaS nanopore displays a higher signal-to-noise ratio (SNR, 21 versus 11 for SiS nanopore) in DNA sensing, although the SNR can be further improved using thinner membranes and smaller pores. The SaS nanopore presents a simple platform in both fabrication and structure that is particularly suitable for low-noise and high-speed molecular diagnostics.

## 1. Introduction

Solid-state nanopores have attracted a lot of interest as a potentially high-speed, portable and low-cost solution for detecting a variety of biomolecules, such as proteins,^[1]^ RNA ^[2, 3]^ and DNA,^[4]^ and studying molecular interactions.^[5, 6]^ However, fundamental limitations in design and manufacturing of low-noise nanopore devices still remain. Currently, a major challenge in prevalent silicon (Si) supported (SiS) solid-state nanopore sensing is associated with a large device capacitance resulted from the Si conductivity. This capacitance introduces a large noise current that becomes particularly dreadful at high recording frequency, thus causing serious reading errors. To mitigate the noise, molecular sensing is often performed at a low bandwidth (*e.g.* 1 kHz to 10 kHz), despite the availability of low-noise, low-current amplifiers operating at much higher (100 kHz and 1 MHz) bandwidth.^[7–9]^ Yet, demoting recording bandwidth seriously limits the signal temporal resolution to ~100 microseconds, in face of the fact that the typical translocation time of a single DNA base pair lies in the range 10-1,000 nanoseconds.^[10]^ To resolve the signals with a high fidelity, a number of methods have been proposed to slow down the DNA translocation speed by reducing its mobility^[11, 12]^ or the effective external DNA-driving force.^[11, 13]^ However, resorting to these methods would introduce high complexity in experiments and decrease the signal-collecting throughput.

An alternative solution without sacrificing the sensor performance is to reduce the noise from the sensing system and the nanopore device (more details in supporting note 1). For instance, a recent demonstration using a customized amplifier and a small-capacitance chip has demonstrated high-speed response of sub-microsecond temporal resolution.^[14]^ Indeed, the Si chip capacitance can be as large as nano-farad range if not carefully engineered (Figure S1c and Table S1). To minimize the stray capacitance, conventional techniques (Table S2) introduce a thick insulating material at the nanopore vicinity,^[14, 15, 16, 17]^ *e.g.* by selective thinning a thick membrane, dielectric coating at nanopore-surrounding areas, or a combination of the two. However, many fabrication schemes require complex and manual processing, such as thick dielectric deposition, selective membrane thinning, electron beam lithography, silicone/photoresist printing, glass bonding, *etc*, and thus are expensive, slow, difficult to reproduce, and not always available to nanopore users. An alternative is to replace conductive silicon by an insulating material, such as glass.^[8, 9, 18–21]^ However, the amorphous nature of the glass substrate makes it impossible to employ selective and anisotropic wet etching to create uniform membranes, which is a favorable high-throughput and inexpensive fabrication technique that has been very successful in forming cavities and suspending membranes on crystalline Si wafers by alkaline solution etching. The fabrication of suspended membranes on glass would have to address challenges in process fluctuation during bulk wafer etching that can adversely affect membrane dimension and geometry, and usually demands complex fabrication schemes involving multiple steps of lithography, deposition and etching on individual chips. Accordingly, the process can cause problems in low fabrication yield, poor reproducibility, and low throughput, seriously limiting the broad availability of such glass-supported nanopore chips.

In this study, we demonstrate a manufacturable approach to create thin membranes with well-controlled dimension and shape on a crystal sapphire wafer, which completely eliminates the stray capacitance that is dominant for conventional SiS nanopore chips for high-speed low-noise sensing. In our approach, we form sapphire-supported (SaS) nanopore membranes by wet and anisotropic etching of sapphire wafers in concentrated sulfuric and phosphoric acids, similar to alkaline etching of Si. Uniquely, we design a triangular membrane by leveraging the three-fold symmetry of the hexagonal c-plane sapphire lattice, and employ a batch-processing compatible anisotropic sapphire wet etching process to create sapphire chips over a wafer scale. We demonstrate that the membrane dimension is controllable in a wide range from ~200 μm to as small as 5 μm, which theoretically corresponds to pico-Farad level total chip capacitance even considering nanometer-thin membranes needed in high-sensitivity DNA detection. Comparing to a SiS nanopore chip, a SaS nanopore chip with a 100 times larger membrane area still has more than two-order-of-magnitude smaller device capacitance and only about one third of current noise measured over 100 kHz bandwidth. Further, the SaS nanopore outperforms the SiS nanopore in high-frequency detection of DNA molecules, demonstrating twice as high SNR despite of having about twice as large pore diameter and 30% thicker membrane. Clearly, further decreasing the membrane area and thickness and creating smaller nanopores in future studies will greatly improve the detection SNR of SaS nanopores for high-speed molecular diagnostics in a wide range of applications.

## 2. Silicon oxide (SiO_2_) supporting membrane formation

We have devised a new strategy to create suspended dielectric membranes on sapphire by anisotropic wet etching (details in Experimental section). Briefly, we started with cleaning a bare 2-inch c-plane (0001) sapphire wafer (**Figure 1a**) by RCA2 cleaning prior to depositing silicon dioxide (SiO_2_) by plasma-enhanced chemical vapor deposition (PECVD) on both sides (Figure 1b). SiO_2_ is used here for its high-selectivity in masking sapphire etching, experimentally determined by us as ~500:1. This was followed by thermal annealing to release the SiO_2_ stress, which otherwise could result in film crack during high-temperature sapphire etching (Figure S2). Then we patterned one side (cavity side) of the SiO_2_ by photolithography and reactive-ion etching (RIE) into a triangular shaped mask layer (Figure 1c). Subsequently, hot sulfuric acid and phosphoric acid (~300°C estimated from previous reports^[22]^) were used to etch through the sapphire wafer to suspend the SiO_2_ membrane (Figure 1d).

**Figure 1.**
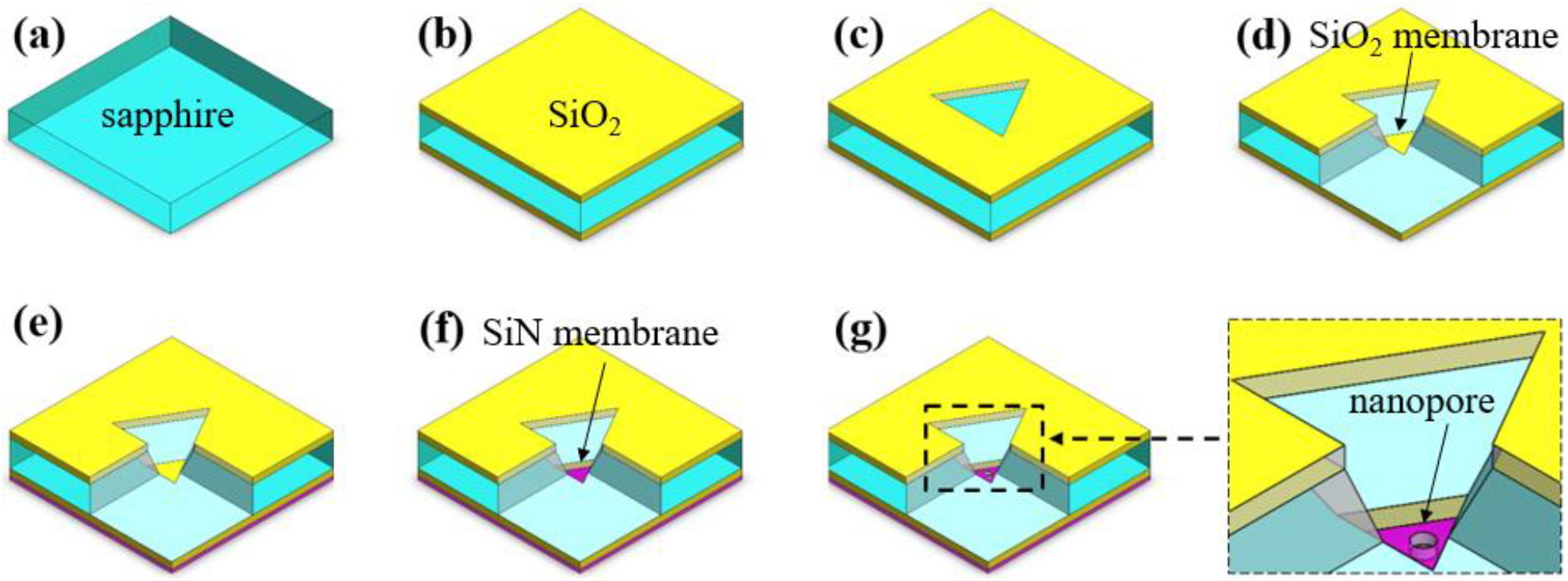
Schematics showing the key steps for creating sapphire-supported nanopores. (a) A 250 μm sapphire wafer is cleaned by solvents and RCA2. (b) A layer of PECVD SiO_2_ is deposited on both sides of the sapphire wafer, followed by thermal annealing. (c) A window is formed in the top SiO_2_ by photolithography and RIE. (d) The sapphire is etched through in hot sulfuric acid and phosphoric acid, forming a suspended SiO_2_ membrane. (e) A thin layer of LPCVD SiN is deposited on the bottom SiO_2_ membrane, and the unintentionally deposited SiN in the cavity is etched by RIE to expose the SiO_2_ membrane in the cavity (not shown). (f) The thin SiN membrane is formed by firstly selectively removing the SiO_2_ membrane in the cavity using hydrofluoric acid and then thinning the SiN using hot phosphoric acid. (g) A nanopore is drilled by transmission electron microscope (TEM) on the SiN membrane. One corner of the chip was hidden in schematic d-g to better show the central etching cavity.

Considering the three-fold symmetric crystal structure of c-plane sapphire wafer, we designed the SiO_2_ etching window as a triangle in order to obtain uniform membrane shape and dimension. The sapphire facet evolution is highly dependent on the alignment of the etching mask to the sapphire crystal, similar to alkaline etching of Si, but more complex given its hexagonal lattice nature ^[23]^. We studied the geometry evolution of the SiO_2_ membrane by rotating the SiO_2_ membrane relative to the sapphire crystal (Figure S3). In another word, we kept the triangular mask dimension the same but changed its alignment angle to the sapphire flat (A-plane), denoted as window-to-flat angle α, and indeed found intriguing formation of membranes. For example, two different sets of triangular membranes were formed when 0 < α < 20° and 40° < α < 60°, with a rotational angle offset between the two at ~30°. In contrast, complex polygon membranes with up to nine sides emerged when 20° <α< 40°, where six of the sides were parallel to the sides of the above-mentioned two triangular membranes. Additionally, the membrane area was also found sensitive to α, yielding an area of more than three orders of magnitude larger when α~30° compared to α~0°. Here we believe the facet evolution is related to the etching rate differences between different sapphire crystal planes. Given that the M- and A-planes have very slow etching rates and are perpendicular to the c-plane, they are believed to be less relevant in the observed cavity formation. We suspect that the R- and N-planes of the sapphire crystals are most relevant,^[24]^ and their competition could result in the angle-dependent evolution into membranes in triangles or nonagon. Drastically different from the triangular design, square window design, which is used in Si etching for its cubic lattice structure, only produced irregular and complex membranes that are much more difficult to control (Figure S4).

Here we chose a designed alignment angle of α~0° and performed theoretical calculation to estimate the relationship between the membrane and the mask dimensions (details in supporting note 2). We determined that the membrane triangle length *L*_2_ could be simply engineered by the mask triangle length *L*_1_ following 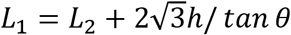 (**Figure 2a**), where ℎ is the sapphire wafer thickness and *θ* is an effective angle between the exposed facets in the cavity and sapphire c-plane that can be empirically determined.

**Figure 2.**
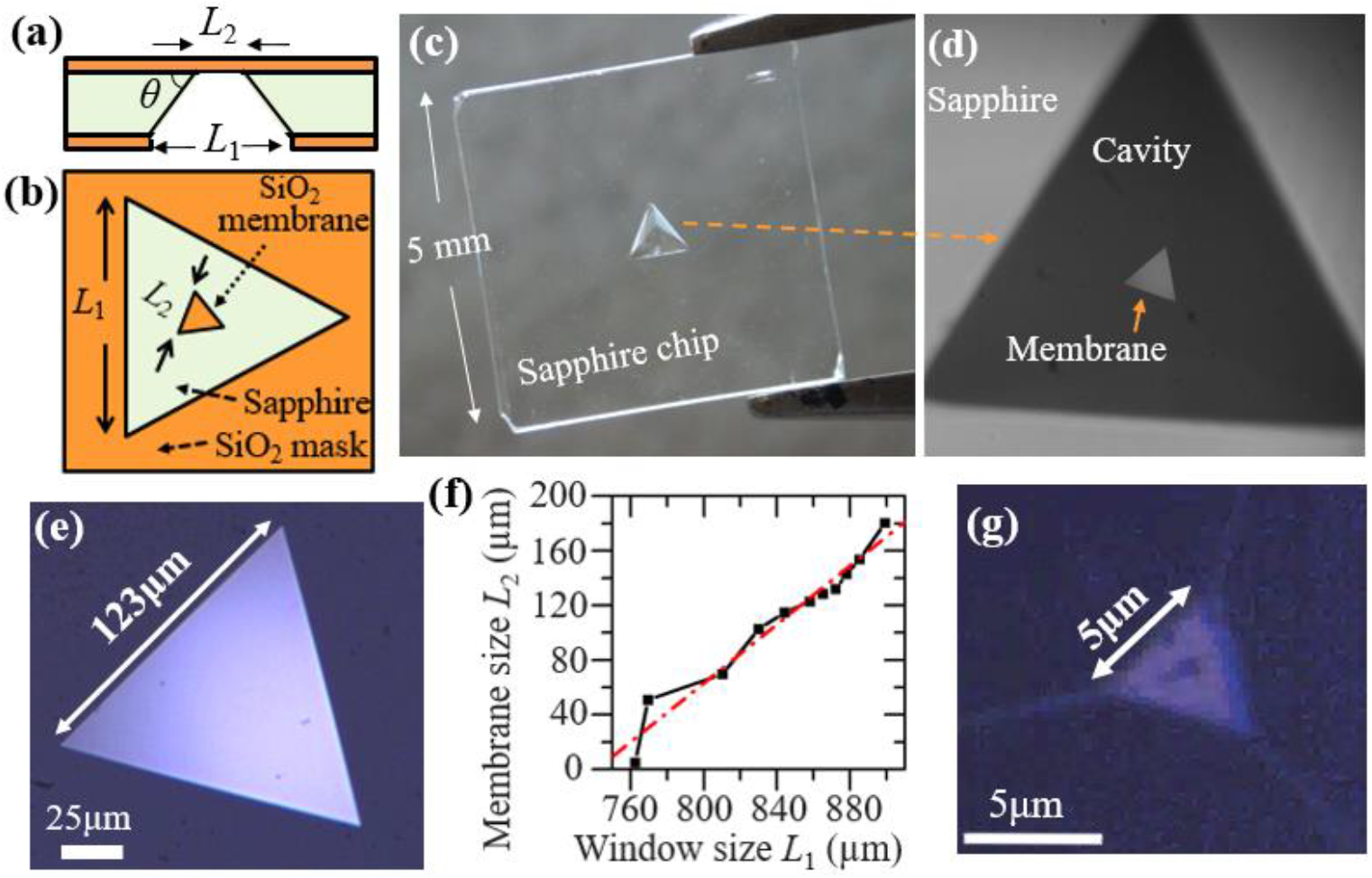
Experimental demonstration of SiO_2_ membrane formation after sapphire etching. (a) Side-view schematic of the chip. L_1_ and L_2_ are the window and the membrane side lengths, respectively. θ is defined as the effective facet angle after etching. (b) Top-view schematic of the chip. (c) An optical image of a 5 mm by 5 mm sapphire chip with intact SiO_2_ membrane. (d) Optical image showing both the triangular window and the SiO_2_ membrane. (e) Optical image of a representative triangular SiO_2_ membrane (L_2_ = 123 μm). (f) Quasi-linear relation between L2 and L1. (g) Optical image of a representative small SiO_2_ membrane (5 μm).

We also intentionally included rectangular dicing marks surrounding the cavity etching windows during lithography, creating trenches in sapphire after acid etching that allowed us to hand-dice sapphire into 5 mm by 5 mm square chips (Figure 2c), which would otherwise be very challenging given the hexagonal lattice of sapphire. This 5 mm chip size was designed to fit into our fluidic jig and transmission electron microscopy (TEM) holder for nanopore drilling and electrical characterization. The final obtained SiO_2_ membrane on sapphire was 3 μm thick and intact during the etching and chip dicing process (Figure 2d-e). The SiO_2_ thickness was only reduced slightly from the original 3.5 μm while masking the etching of 250 μm sapphire, indicating an ultra-high etching selectivity of ~500:1. The SiO_2_ membrane size *L*_2_ was also found tunable in a wide range from 5 to 200 μm (Figure 2f-g, more images in Figure S5a). The 5 μm membrane corresponds to a theoretical pico-farad chip capacitance even for nanometer-thin membranes, *e.g.* <2 pF for a hypothetical 2 nm thick two-dimensional (2D) material membrane^[25]^ (more details in Table S3), which are highly desired for high-SNR^[3, 26]^ DNA detection. We further fitted the correlation between *L*_1_ and *L*_2_ using our theoretical model, and determined an effective facet angle θ~ 50° (Figure S5d). This experiment proved that it was possible to control and create ultrasmall membranes for functional sapphire chips. It was also intriguing to notice the complex sapphire facets from scanning-electron microscope (SEM) image of the formed cavity (Figure S6a), attributed to the complex crystal structure of sapphire and particularly possibly due to the competition between R- and N-planes of the sapphire crystals. These studies serve to establish a new fabrication and design strategy that create unique triangular membrane devices through bulk anisotropic etching of sapphire wafers, which is crucial to the implementation of SaS nanopore chips. Additionally, our established integration scheme will inspire translation of conventional Si based micromachining strategies and microelectromechanical systems (MEMS) designs into new sapphire based micro-devices.^[27]^

## 3. SiN thin membrane formation

Using the triangular SiO_2_ membranes formed by sapphire etching, we have developed a process to create thin silicon nitride (SiN) membranes suitable for nanopore formation and DNA sensing.^[3]^ Briefly, we deposited low-stress SiN film on the suspended SiO_2_ membranes by low-pressure chemical vapor deposition (LPCVD), and then removed the SiO_2_ film within the triangular aperture *via* selective dry etching and HF based wet etching from the cavity side (Figure 1f). Using the SiN film instead of the remaining SiO_2_ mask layer as the membrane material allows us to precisely control the membrane thickness and largely eliminates high compressive stress from the SiO_2_ layer that negatively affects the membrane integrity. Starting from a 320 nm-thick SiN film that was structurally robust, we evaluated both RIE and wet etching in hot phosphoric acid to thin down the SiN membrane to desired thickness. We found that RIE could cause non-uniformity (Figure S7a) and might damage the membrane, causing current leakage, as shown by current-voltage (*IV*) characteristics using one molar potassium chloride solution (1M KCl) (Figure S7b). In contrast, wet etching in hot phosphoric acid yielded uniform SiN membrane (Figure S7c and Figure S8b) without current leakage (Figure S7d), thus preferable for the DNA sensing test. Finally, a nanopore was drilled in the SiN membranes on the sapphire chip (**Figure 3a-b**) and a float-zone Si chip (SiMPore Inc., Figure S9) by TEM (Figure 1g) for electrical characterization and DNA sensing test.

**Figure 3.**
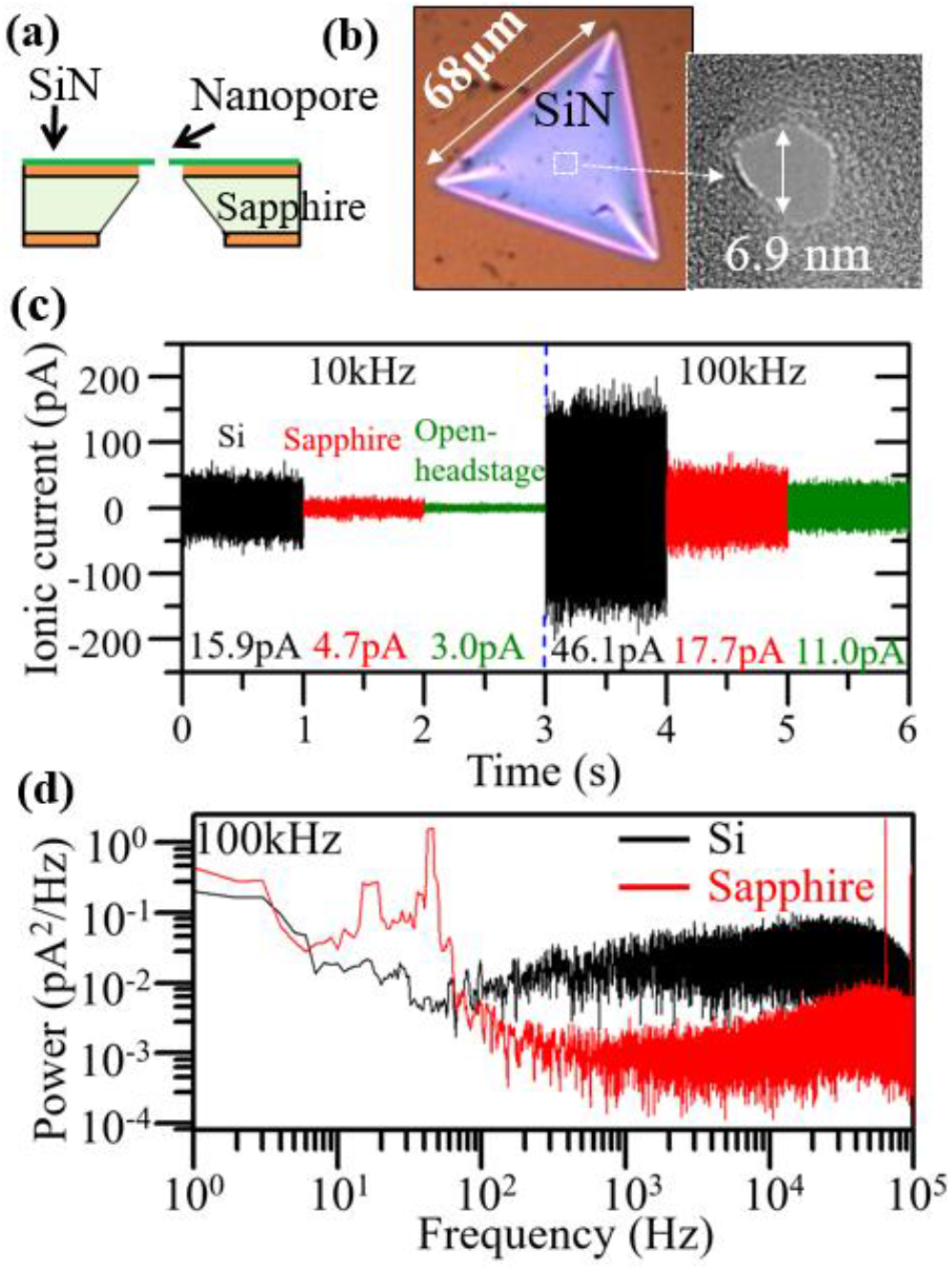
Ionic current noise analysis of a SaS nanopore and a SiS nanopore. (a) A schematic of the measured SaS nanopore chip. (b) An optical image of the SiN membrane of the SaS nanopore chip and a TEM image of the drilled nanopore. (c) The ionic current noise for the SiS nanopore (black traces), the SaS nanopore (red traces), and the open-headstage state (green traces) under 10 kHz (left three traces) and 100 kHz (right traces) low-pass filter respectively. The two chips were both measured under 50 mV voltage. The RMS ionic current values are also analyzed and marked for each measurement. (d) Power spectra of the current noise of the SaS nanopore and the SiS nanopore versus frequency under 100 kHz low-pass filter. The two chips were both measured under 50 mV voltage.

## 4. Noise characterization

First we experimentally characterized the device capacitance of the SaS and SiS nanopore chips. Noticeably, the SaS nanopore chip had a 100 times larger membrane area (68 μm triangular side length, or ~2000 μm^2^ in area) than the SiS nanopore chip (4.2 × 4.7 μm square, or ~20 μm^2^ in area) and slightly thicker SiN (30 nm for sapphire and ~23 nm for Si). Following 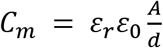, where *C*_*m*_ is the membrane capacitance, *ε*_*r*_ is the relative permittivity of SiN, *ε*_0_ is the vacuum permittivity, *A* is the membrane area and *d* is the membrane thickness, we calculated the SaS nanopore membrane capacitance as 3.8 pF, more than 70 times bigger than that of the SiS nanopore chips (0.05 pF). However, the SaS chip was experimentally found to have a much smaller total capacitance (~10 pF) compared to the SiS chip (~1.3 nF) using the Clampex software (Molecular Devices, LLC). Clearly, considering SaS and SiS nanopores that both have only the simplest membrane structure, it is clear that insulating sapphire successfully eliminated the dominant capacitance resulting from Si substrate conductivity, thus appealing to low-noise measurement.

We also compared the SaS chip capacitance to that of some representative low-noise SiS chips and glass-supported chips (Table S2). The capacitance of SiS membranes^[16, 17]^ can be engineered to <10 pF by the combination of two widely adopted approaches, namely reducing the membrane area and depositing thick insulating layers surrounding the membranes. An ultrasmall nanometer-scale membrane is usually implemented by electron-beam lithography (EBL) followed by precisely timed etching, which are complex, expensive in training and fabrication, and inherently susceptible to processing fluctuations. Insulating layers can be introduced by aligned glass bonding, film deposition followed by aligned lithography and etching, and even silicone painting. However, the required sophisticated instrument, extensive training, and manual operation (alignment or painting) make the fabrication of low-noise SiS membrane chips costly, time-consuming, and low-yield. Glass-supported nanopore chips^[9, 18–20, 28]^ have been demonstrated to have shown a reduced capacitance as small as ~70 pF^[18]^ with 5 μm diameter membrane or even <10 pF^[19, 20, 28]^ with as small as 100-300 nm-diameter membrane. In comparison, our SaS chip with a relatively large membrane (68 μm triangular side length) measured only ~10 pF capacitance - a value comparable to that of the state-of-art glass-supported nanopore chips. Additionally, the amorphous glass wafer, typically a few hundreds of micrometers in thickness, is usually etched in hydrofluoric acids (HF) to minimize manufacturing cost for membrane formation. However, the deep isotropic etching in HF would unenviably cause serious lateral undercut, making it very challenging to precisely control the membrane dimension for capacitance minimization. Better membrane dimension control is possible by introducing RIE etching, but it would require extra processing at single-wafer or single-chip level and drastically lower the throughput and increase the manufacturing cost. Recently, it has shown that pulsed femto-second laser ablation, in combination with LPVCD of SiN and chemical wet etching, can be used to form nanopores in glass with a device capacitance as small as ~2 pF^[20, 29]^ The results are very encouraging, yet it remains to be seen how process fluctuation in laser focusing, laser power, and etching would affect the precision in controlling the membrane uniformity (currently reported variation from 5 to 40 μm) and the fabrication throughput.

We further analyzed the ionic current noise for the SaS nanopore, the SiS nanopore and the open-headstage system (Axopatch 200B) under 10 kHz and 100 kHz low-pass filter (Figure 3c). The root-mean-square (RMS) of the measured current of the SaS nanopore chip is ~5 and 18 pA using 10 and 100 kHz filters, only slightly higher than the open-headstage system RMS noise (3 and 11 pA) but much better than those from our SiS nanopore (~16 and 46 pA). In comparison, the best reported silicone-painted SiS chips^[16]^ that utilized a locally thinned membrane (0.25 μm^2^ area and 10-15 nm thick in the center) produced ~7 and ~13 pA noise current at 10kHz and 100kHz, measured by an optimally designed amplifier that greatly outperforms Axopatch 200B at a high recording frequency (>10kHz). Additionally, we also compiled the reported noise current from glass-supported nanopore chips (Table S2). For example, one of the best glass chips with nano-membrane (*e.g.* 100 nm diameter, 0.008 μm^2^ in area, and 5-10 pF) measured ~4 and ~13 pA noise current at 10 kHz and 100 kHz, respectively.^[28]^ Glass chips with micro-membranes (25 μm^2^ and 70 pF,^[18]^ and 314 μm^2^, ~2 pF)^[20]^ measured ~13 and ~19 pA noise current at 10 kHz bandwidth, about 3-4 times larger than our SaS chip. The above comparison convincingly shows that our SaS chips are successful in suppressing the noise current and completely comparable to the best reported Si- and glass-supported nanopore chips.

Additionally, our analysis of the power spectral density (PSD) (Figure 3d) further demonstrated that the SaS nanopores outperformed the SiS chips, particularly at high bandwidth (*e.g.* >10 kHz) due to the significantly reduced device capacitance. In the moderate frequency range (*e.g.*100 Hz to 10 kHz), the noise power of SaS nanopore was about one order of magnitude smaller (~10^−3^ pA^2^/Hz) than that of our measured SiS nanopore (~10^−2^ pA^2^/Hz) and one order of magnitude smaller or comparable to the glass chips and low-noise SiS chips (10^−2^~10^−3^ pA^2^/Hz)^[16–18, 20, 28]^ (Table S2), partly attributed to lower dielectric noise and Johnson noise. We note that sapphire has a very small dissipation factor *D* (~10^−5^), two to five orders of magnitude smaller than that of typical borosilicate glass (10^−3^ to 10^−2^) and Si (1-100) and comparable to that of high-purity fused silica (~10^−6^).^[30]^ Such a small dissipation factor, together with its small device capacitance, is favorable for minimizing noise related to dielectric loss (dielectric noise^[31]^ *S*_*D*_ ∝ *DC*_*chip*_*f*, *C*_*chip*_ is nanopore chip capacitance and *f* is the frequency). Additionally, the high resistivity of sapphire (>10^14^ Ω·cm) also served to minimize resistance-related Johnson noise. At very low frequency range (<100 Hz), the noise power of the SaS nanopore was about 10^−1^ pA^2^/Hz, one order higher than SiS nanopore (10^−2^ pA^2^/Hz), which could be attributed to the flicker noise and could be further improved by surface modification.^[32]^ From the above analysis, it is again evident that SaS chips are completely comparable to the best available SiS and glass-supported nanopore chips. Yet, our demonstrated batch-processing-compatible design, significantly simplified fabrication process, and comparable price to high-quality glass (fused silica) make sapphire an excellent candidate for low-noise and high-frequency nanopore sensing at a low cost.

## 5. DNA detection

To evaluate the performance in the detection of DNA molecules by our SaS nanopore, 1kbp ds-DNA translocation events were measured under 100 kHz (**Figure 4**) and 10 kHz (Figure S10) low-pass filter for both the SaS and the SiS nanopore under 50 mV, 100 mV and 150 mV bias. Studying representative ionic current traces of 1kbp dsDNA (Figure 4b) for both SiS and SaS nanopores, we note that the DNA signals collected by SiS nanopore were more irregular, particularly at lower bias voltages. These irregular signals, together with the high baseline noise, made it very challenging to faithfully distinguish DNA signals from the background. In comparison, the SaS nanopore produced DNA signals with much less distortion or noise at 100 kHz bandwidth that can be easily identified. Additionally, we also show that recording at lower frequencies (such as 10 kHz) would result in serious data loss of the fast DNA signals, thus presenting only longer and in some occasions distorted signals.^[16, 28]^ Clearly, SaS nanopores enable preferable high-speed, high-throughput, and high-fidelity detection of DNA signals.

**Figure 4.**
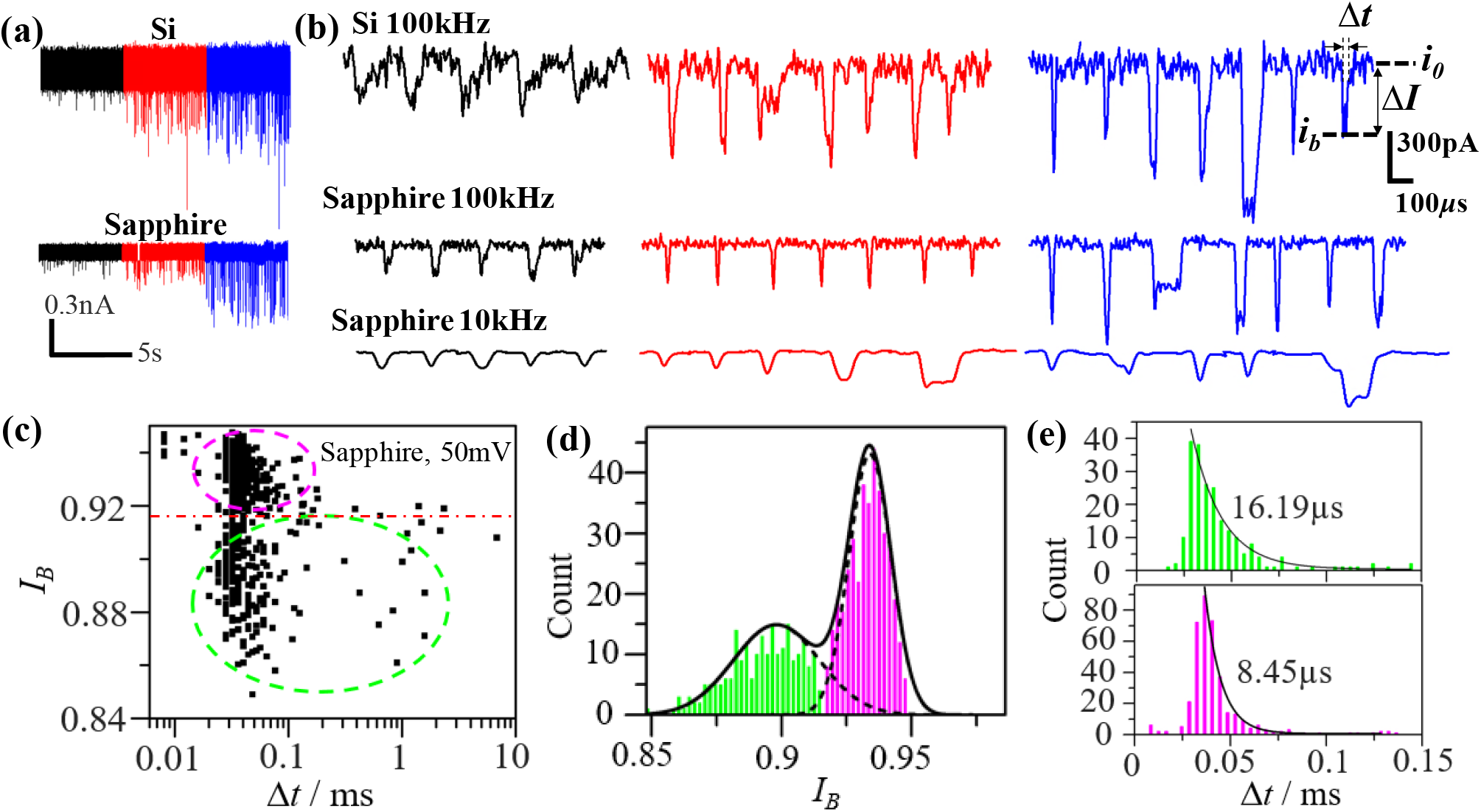
Analysis of 1k bp dsDNA translocation events for the SaS nanopore (2002 μm^2^ membrane area) and the SiS nanopore (31 μm^2^ membrane area) under 100 kHz filter frequency. (a) The current traces of the DNA translocation events of the SiS nanopore and the SaS nanopore under different voltages (black: 50 mV, red: 100 mV, blue: 150 mV). (b) Representative DNA events for the SiS nanopore and the SaS nanopore at different voltages (black: 50 mV, red: 100 mV, blue: 150 mV) and different recording bandwidth (top two rows: 100 kHz, bottom row: 10 kHz). Δt: event dwelling time; *i_0_*: open-pore current baseline; *i_b_*: block-pore current level; ΔI: blockade current amplitude. (c) Scatter plot of the fractional blockade current I_B_ (=*i_b_*/*i_0_*) versus the dwelling time Δt of all the DNA events from the SaS nanopore under 50 mV. Two distinct populations are separated by the red dashed line as the translocation events (green oval) and the collision events (pink oval). (d) The histograms of I_B_ of the SaS nanopore under 50 mV displaying two distinct peaks corresponding to the translocation events (green bars) and the collision events (pink bars). The solid and dash black lines indicate the fitting by Gaussian function. (e) Histograms of Δt of the segregated events based on two I_B_ populations, fitted by exponential function. The translocation events (top panel) has a longer tail (decay constant 16.19 μs) than the collision events (lower panel, decay constant 8.45 μs).

To study the DNA translocation mechanism, we extracted the DNA signals by OpenNanopore Program.^[33]^ We scatter-plotted the fractional blockade current *I_B_* (=*i_b_/i_0_*) and the dwelling time Δ*t* of all the DNA events from the sapphire chip under 50 mV (Figure 4c). Here *i_b_* is the blocked-pore current and *i_0_* is the open-pore current. The use of *I_B_* allows us to eliminate the impact of bias difference on DNA signal analysis. Two distinct populations were observed (separated by the red dashed line in Figure 4d) and recognized as the translocation events (green oval) and the collision events (pink oval).^[5]^ Further, we analyzed the current blockade distribution and fitted with Gaussian function (Figure 4d), producing two distinct *I_B_* populations attributed to translocation and collisions. We further analyzed the dwelling time Δ*t* of each of the two event populations and fitted with exponential decay function (black lines, Figure 4e). It showed that the translocation events (green, top panel) had a longer tail (decay constant 16.19 μs) than the collision events (decay constant 8.45 μs), consistent with previous studies.^[5]^

We further applied this signal segregation approach to analyze all the DNA signals collected from the SiS and SaS nanopores (**Figure 5a-d**). By scatter-plotting the normalized DNA blockade signal (1-*I*_B_ = Δ*I*/*i_0_*) and marking the normalized current noise (*I_RMS_*/*i_0_*, dash-dot lines) at each bias voltage (black: 50 mV, red: 100 mV, blue: 150 mV, Figure 5e-f), we could investigate the SNR (defined here as 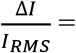 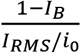 of the true DNA translation signals. The short solid lines represented the average DNA signals (1-*I*_*B*_) determined from the Gaussian distribution of the translocation events (Figure 5b, d). The SaS nanopores produced slightly smaller DNA signal amplitude than SiS nanopores, because of their larger pore size and thicker membrane. However, given the suppressed noise current, the SaS nanopore still evidently outperformed SiS nanopore in SNR. For example, the SaS nanopore has a SNR of 21 at 150 mV bias, almost twice as good as the SiS nanopore.

**Figure 5.**
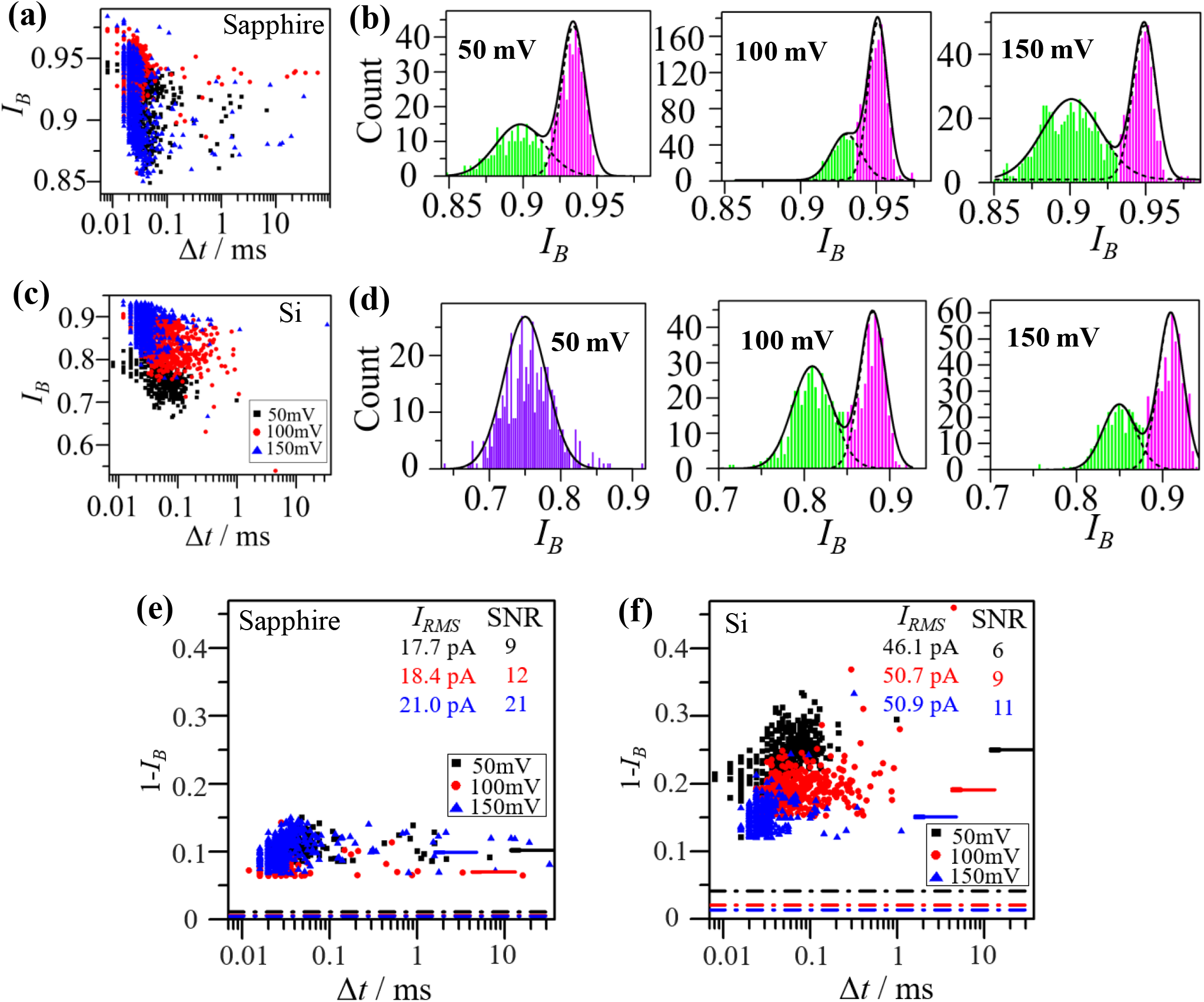
Signal-to-noise ratio (SNR) comparison between the SaS nanopore and the SiS nanopore under 100 kHz filter frequency. (a) Scatter plot of the fractional blockade current I_B_ (=*i_b_*/*i_0_*) versus the dwelling time Δt of all the DNA events from the SaS nanopore under different bias voltages from 50 mV to 150 mV. (b) The histograms of I_B_ of the SaS nanopore. Two distinct peaks are observed and fitted by Gaussian function, corresponding to the translocation events (green bars) and the collision events (pink bars). (c) Scatter plot of the fractional blockade current I_B_ (=*i_b_*/*i_0_*) versus the dwelling time Δt of all the DNA events from the SiS nanopore. (d) The histograms of I_B_ of the SiS nanopore. Two distinct peaks are observed for 100 mV and 150 mV biases and fitted by Gaussian function, corresponding to the translocation events (green bars) and the collision events (pink bars). The signals at 50 mV bias displayed only one obvious peak and not further segregated. (e-f) Scatter plot of 1-I_B_ (=ΔI/*i_0_*) versus the dwelling time Δt of all the DNA translocation events (collision events removed) from the SaS nanopore (e) and SiS nanopore (f). The dashed lines at the bottom are the values of I_RMS_/*i_0_*, in which IRMS is the root-mean-square noise at open-pore state. The short solid lines are the peak values of (1-I_B_) in the Gaussian distribution of the translocation events in (b) and (d). The error bars of the distribution are added at the left edge of each short solid line. The SNR for each bias voltage is determined by the ratio between the values of the DNA signals, indicated by the short solid lines, and their corresponding noises, represented by the dashed lines of the same color. The values of SNR are also marked in the figures. DNA data are represented by black, red and blue dots in figure a, c, e, and f for the collecting bias voltages as 50 mV, 100 mV, and 150 mV.

We further attempted to detect short single-stranded (ss) DNA molecules using SaS nanopores (**Figure 6**). Ionic current traces of Poly(A)_40_ ssDNA translocation events were recorded under 100 kHz low-pass filter with the voltages from 100 mV to 150 mV. We performed the same analysis to investigate the SNR of this ssDNA (Figure 6b and Figure S11), and obtained a SNR of ~6 for both 100 mV and 150 mV bias voltages. This provided evidence that the SaS nanopores can detect a wide range of biomolecules of different sizes. We expect the SNR can be remarkably enhanced by using thinner membrane thickness and small nanopore in future studies.

**Figure 6.**
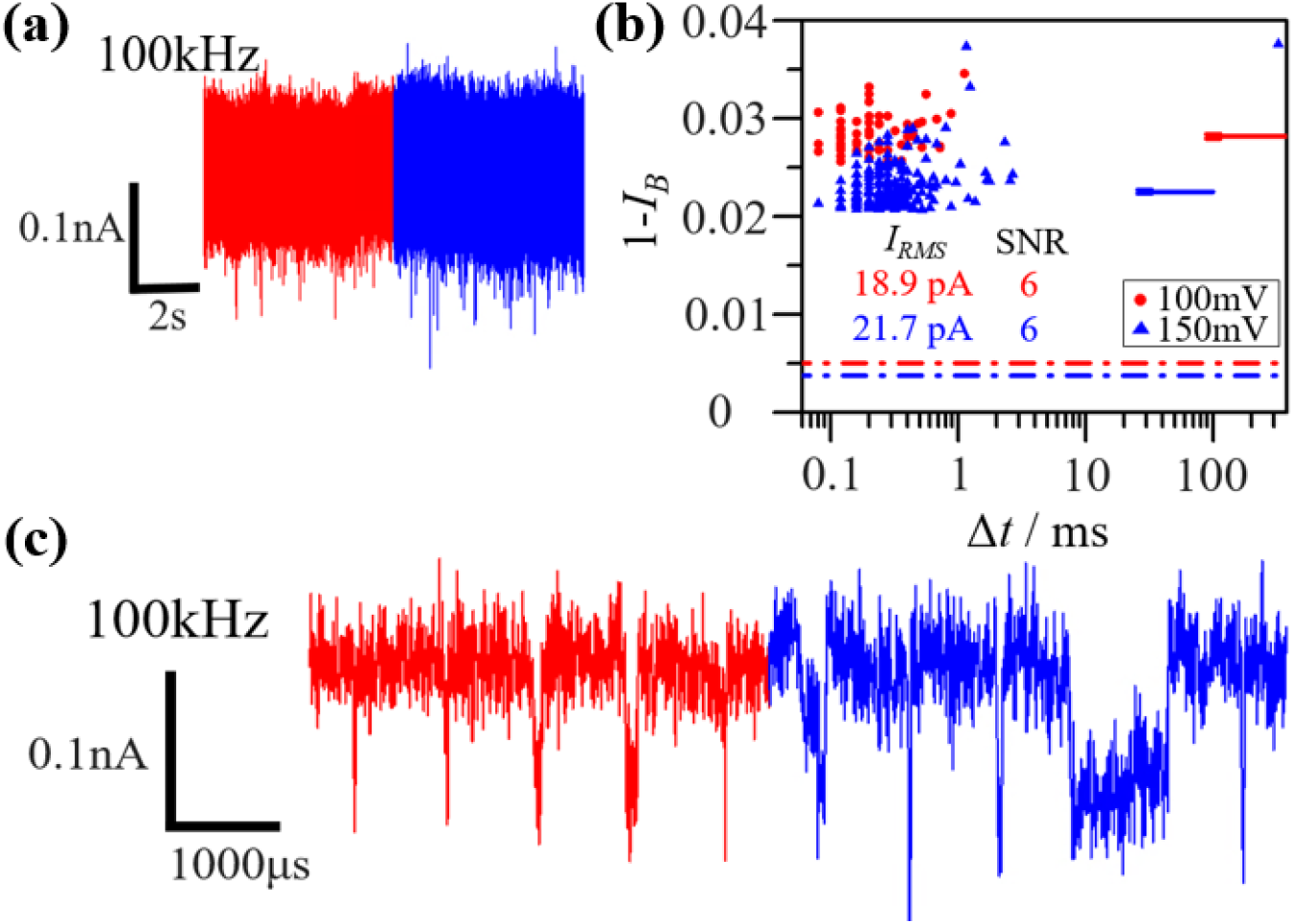
Analysis of Poly(A)_40_ single-stranded (ss) DNA translocation events for the SaS nanopore under 100 kHz filter frequency. (a) The current trace of the DNA translocation events under 100 kHz filter frequency. (b) Scatter plot of 1-I_B_ (=ΔI/*i_0_*) versus the dwelling time Δt of all the DNA translocation events (collision events removed). The dashed lines at the bottom are the values of I_RMS_/*i_0_*, in which IRMS is the root-mean-square noise at open-pore state. The short solid lines are the peak values of (1-I_B_) in the Gaussian distribution of the translocation events. The error bars of the distribution are added at the left edge of each short solid line. The SNR is given by the ratio of the DNA signal (short solid lines) and the noise (dashed lines) for each tested voltage. (c) Representative DNA events under 100 kHz filter frequency. Here the signals are indicated by red and blue for bias voltages at 100 mV and 150 mV, respectively.

## 6. Conclusion and outlook

In conclusion, we demonstrate a novel design and manufacturable approach to create SaS nanopores featuring triangular membranes with well-controlled dimensions and shapes. Completely eliminating the stray capacitance, the SaS nanopores convincingly produced two-order-of-magnitude smaller device capacitance (10 pF) compared to a measured SiS nanopore (~1.3 nF) despite having a 100 times larger membrane area. Accordingly, the SaS nanopores generated ~5 times smaller RMS ionic current noise than the SiS nanopore at 100 kHz bandwidth, and resulted in high-fidelity DNA sensing with a twice higher SNR while having a larger nanopore size and thicker SiN membrane. By analyzing the device capacitance, noise current, and power density spectra of the SaS nanopore and comparing to the best reported SiS and glass-supported nanopores, we found our nanopore chips were comparable to the best available low-noise sensors. Further optimization in reducing the membrane area and membrane thickness is expected to further decrease the device capacitance and noise, increase the sensitivity, and boost the SNR. Particularly, integration with ultrathin 2D materials^[25]^ (such as MoS_2_, WS_2_, h-BN, *etc.*) as the membrane is viewed as a promising approach to create highly sensitive nanopores. It is even possible to eventually pursue an all-sapphire nanopore device platform, where the membrane is also made of ultrathin sapphire that can facilitate the Van der Waals epitaxial growth of 2D materials for scalable 2D membrane deposition.

This low-capacitance SaS membrane is very favorable to future integration with scalable nanopore formation technologies that require ultrafast feedback from voltage/current signals, such as dielectric breakdown^[34]^ and laser based nanopore drilling.^[35, 36]^ Additionally, this structurally ultra-simple and optically transparent platform makes it particularly attractive for coupling optical spectroscopy and fluorescent imaging with electrical signal readout.^[36]^ which has broad applications in studying the complex DNA conformational changes and dynamic interaction with nanopore surface and improving electrical signal deconvolution. The SaS nanopore platform will also find use in detection of a variety of biomolecules besides DNA, such as RNA, protein, extracellular vesicles, *etc*. For example, the low-capacitance feature is particularly beneficial to minimizing the RC delay time while implementing automatic recapturing in the studies of the mechanical deformation of extracellular vesicles^[37]^ and the protein-protein interactions.^[38]^ Beyond nanopores, our batch-processing compatible and potentially cost-effective manufacturing of the SaS membrane architecture, together with the high mechanical strength, chemical resistivity, high temperature stability, and high optical transparency of sapphire, may serve to establish a new fabrication and design strategy in bulk micromachining of sapphire wafers to broaden the applications in MEMS designs and optoelectronic devices.^[39]^

## 7. Experimental Section

### (1) SaS nanopore membrane fabrication

Firstly, a 250 μm thick 2-inch c-plane sapphire wafer (Precision Micro-Optics Inc.) was treated by RCA2 cleaning (deionized water: 27% hydrochloric acid: 30% hydrogen peroxide = 6: 1: 1, 70 °C) for 15 min followed by 3.5 μm PECVD SiO_2_ deposition (Oxford PECVD, 350 °C, 20 W, 1000 mTorr, SiH_4_ 170 sccm, N_2_O 710 sccm, deposition rate: 68 nm/min) on both sides. Then the wafer was brought in a furnace for thermal annealing (400 ° C, 2 hrs, air ambient) to release the stress in SiO_2_ film, followed by photolithography (Heidelberg Instruments μPG 101 laser writer, 600 nm AZ 1505 photoresist) and RIE (PlasmaTherm 790 RIE Fluorine, 250 W bias, 40 mTorr, CHF_3_ 40 sccm, O_2_ 3 sccm, etching rate: 46 nm/min) etching on SiO_2_ to form a triangular etching window.

Next, hot sulfuric acid and phosphoric acid (3:1, hot plate setting 540 °C, solution temperature ~300°C) were used to etch through the sapphire wafer (etching rate: 12 μm/hr) and suspend the SiO_2_ membrane. To ensure the safety of handling hot and concentrated acids, we custom-designed a quartz glassware setup suitable for high-temperature acid-based sapphire etching process. We intentionally placed the sapphire wafer vertically in a 2-inch glass boat in the etching container to minimize possible damage to the membrane by the boiling acids (Figure S12). After the acid was added into the quartz glassware, we loaded the 2-inch glass boat with the wafer into the quartz glassware, and installed a clamp seal and a condenser column to minimize acid vapor leakage. Finally we raised up the temperature of the hot plate to 540 °C (100-200 °C/min ramping rate) to start the etching. The etching rate was chosen to be relatively slow to minimize wafer breakage during etching in our customized container, but further increasing the solution temperature could exponentially increase the etching rate and allow a larger throughput. We note that commercially available etching tanks are available that can allow automated etching with more precise control in temperature and batch processing.

Following sapphire etching, the SiO_2_ membrane was thinned down by RIE (PlasmaTherm 790 RIE Fluorine, 250 W bias, 40 mTorr, CHF_3_ 40 sccm, O_2_ 3 sccm, etching rate: 46 nm/min) to 1.45 μm, and a layer of SiN (320 nm) was deposited onto the SiO_2_ membrane by LPCVD (Tystar TYTAN 4600, 250 mTorr, DCS flow 25 sccm, NH_3_ flow 75 sccm, 750 °C, deposition rate: 6 nm/min). SiN unintentionally deposited in the back cavity of the chip was removed by a RIE etching step (PlasmaLab 80 Fluorine, 100 W bias, 100 mTorr, CF_4_ 50 sccm, O_2_ 2 sccm, etching rate: 61 nm/min). Then hydrofluoric acid (8%) was used to remove the SiO_2_ layer to suspend the SiN layer (90 nm/min). The final SiN membrane was thinned down by hot 85% phosphoric acid (~130°C estimated from previous reports,^[40]^ etching rate: ~2.5 nm/min) to desired thickness.

### (2) SiS nanopore membrane fabrication

The SiS nanopore membranes were purchased from SiMPore Inc. The chips were made from 100 mm diameter, 200 μm thick, float-zone Si wafer (resistivity of 1-10 Ω∙cm) with ~100 nm thermal SiO_2_ and ~20 nm LPCVD SiN films, where the thermal SiO_2_ from the cavity side was removed to produce an array of 4-5 μm diameter, suspended SiN membranes. The SiO_2_ and SiN film thicknesses were confirmed by M-2000 ellipsometer (J.A. Woollam Co.) as 99 nm and 23 nm by us.

### (3) Thickness characterization on the small membranes

The thicknesses of membranes were measured by Filmetrics F40 (Filmetrics Inc.), which has the capability to measure small area and is based on the reflectance and the refractive index of the measured material. For the LPCVD SiN membranes, the refractive index was first fitted using the same-batch LPCVD SiN deposited on Si by Woollam Spectroscopic Ellipsometer (J.A. Woollam Co.). Then the refractive index list was exported to Filmetrics F40 to measure the thickness of the SiN suspended membrane (film stack: air-SiN-air). A well-fitting curve of the central region of the triangular membrane was shown in Figure S8a.

### (4) Nanopore drilling

The nanopore was drilled by JEOL 2010F TEM. The 5 mm by 5 mm nanopore chip was placed in a customized 5 mm TEM sample holder. The largest condenser aperture and spot size 1 were used for maximum beam current output. After the alignment was finished, the imaging magnification was increased to 1.5M (maximum). The beam spot was spread to 3 inch and held for 5-15 min for stabilization. If the beam spot drifted, the focus needed to be re-adjusted under 250K magnification and the stabilization needed to be re-monitored under 1.5M magnification. Once the beam got stabilized, the 3-inch beam spot was reduced to ~7 mm and the condenser astigmatism was quickly adjusted to make the spot as round as possible. At this stage, from the eyepiece, the material being bombarded could be observed. Once it was clear, a successful drilling was identified. Under the condition of 7.01 kV anode A2 (focusing anode), 3.22 kV anode A1 (extraction anode) and 30 nm SiN membrane, it took 75-90 sec to drill through the membrane.

### (5) Noise characterization, DNA preparation and DNA sensing

The TEM-drilled nanopore chip was treated with UV ozone cleaner (ProCleaner™, BioForce Nanosciences Inc.) for 15 min to improve the hydrophilicity of the surface and mounted into a customized flow cell (Figure S13). Then a solution of 1:1 mixed ethanol and DI water was injected into the flow cell to wet the chip for 30 min. The solution was subsequently flushed away by injection of DI water. Next, 100 mM KCl was injected into the flow cell to test the current-voltage (IV) curve using Axopatch 200B amplifier and Digidata 1440A digitizer (Molecular Devices, LLC.), and then 1M KCl solution was injected to characterize the device current. To do DNA sensing, the 1kbp as-ordered dsDNA (Thermo Scientific NoLimits, Thermo Fisher Scientific Inc.) was diluted using 1M KCl to 5 ng/μL or the Poly(A)40 ssDNA (Standard DNA oligonucleotides, Thermo Fisher Scientific Inc.) was diluted using 1M KCl to 50nM, and stirred using a vortex mixer. Finally, the DNA solution was injected into the flow cell to collect DNA signals under 10 kHz and 100 kHz low-pass filter at 50, 100 and 150 mV using Axopatch 200B amplifier and Digidata 1440A digitizer (Molecular Devices, LLC.). The flow cell was kept in a customized Faraday cage on an anti-vibration table (Nexus Breadboard, Thor labs) to isolate the environment noise during measurement.

### (6) DNA signal collection and analysis

After the injection of the DNA solution, once the external voltage was applied, DNA signal could be observed from the Clampex software. The DNA signals were recorded for sufficient time at each voltage (50, 100, 150 mV) and each frequency (10 and 100 kHz) to ensure a relatively large data set for analysis. The collected DNA signals were analyzed by OpenNanopore program.^[33]^ Firstly we edited a MATLAB program to convert all the .abf files to .mat files in a batch. Then these .mat files were imported to OpenNanopore program to generate the dwelling time and blockade current amplitude data of each DNA signal for subsequent analysis.

## Supporting Information

Supporting Information is available from the Wiley Online Library or from the author.

## Acknowledgments

This work is partially supported by the Arizona State University (ASU) startup funds to Prof. Chao Wang and National Science Foundation under award no. 1711412, 1809997, 1838443, 1847324 and 2020464. We thank Dr. G. Stolovitzky at IBM, Prof. A. Meller at Technion - Israel Institute of Technology, Dr. Y. Astier and Dr. J. Topolancik from Roche Sequencing Solutions, Dr. S. Lindsay at ASU, and Dr. P. Pang (currently with Roswell Biotechnologies) at ASU for fruitful discussions.

