## Supporting information for "Sapphire-Supported Nanopores for Low-Noise DNA Sensing"

**Supporting note 1**

The power spectral density (PSD) of a solid-state nanopore can be described as[1]

$$S = a_1 \frac{1}{f^{\beta}} + a_2 + a_3 f + a_4 f^2 \quad (1)$$

where $f$ is frequency and $a_i (i = 1,2,3,4)$ are coefficients. $S$ consists of the low-frequency flicker noise $\frac{a_1}{f^{\beta}}$ (1< $\beta$ <2),[2] Johnson noise (white thermal noise) $a_2$, dielectric noise[3] $a_3 f$, and capacitive noise[4] $a_4 f^2$. The Johnson noise is correlated to the thermal fluctuations of the charge carriers ($a_2 \propto \frac{T}{R}$, $T$: temperature, $R$: nanopore resistance).[5] The dielectric noise is related to the dielectric loss which originates from the existence of charge carriers in the dielectric material ($a_3 \propto DC_{chip}$, $D$: dissipation factor, $C_{chip}$: nanopore chip capacitance).[5] The noise current at high frequencies (*e.g.* >10 kHz) is mainly contributed by the capacitive noise $a_4 f^2$ (Figure S1b). This noise is proportional to total input capacitance $C_{total}$ with a PSD growing with the square of frequency - $S_{amp} = (2\pi f C_{total} v_n)^2$, where $v_n$ is the voltage noise density of the input equivalent voltage thermal noise of the input amplifier.[6] And in this high-frequency sensing regime, $I_{RMS}(B) = \frac{2\pi}{\sqrt{3}} B^{3/2} C_{total} v_n \sqrt{S(B)}$.[4, 7]

The total input capacitance is estimated as $C_{total} = C_{sys} + C_{chip}$.[4] in which $C_{sys}$ is the capacitance from the measurement setup. $C_{sys}$ is generally on the order of 10-20 pF and the optimized amplifier design can decrease it down to less than 5pF.[7, 8] $C_{chip}$, which is composed of $C_m + C_s$, can be as large as hundreds of pF or even a few nF. $C_m$ is the membrane capacitance, which becomes negligible for small membranes. However, $C_s$ (stray capacitance), which is expressed as $C_{Si-B1} \parallel (C_{Si-B1} + C_{Si-B3})$ (Figure S1c) for first-order estimation, can be significant due to the significant amount of free carriers in silicon (Si) substrate.[7]

**Supporting note 2**

To estimate the relationship between the membrane and the mask dimensions, we assume $L_2$ (membrane side length) is parallel to $L_1$ (window side length) and the etching follows an effective facet angle $\theta$ (Figure S5c), which is between the exposed facets in the cavity and sapphire c-plane that can be empirically determined (Figure 2a). In Figure S5s, three auxiliary lines are added to connect the six vertexes of the two triangles and two auxiliary lines are shot from the two vertexes of the membrane triangle to be perpendicular to the sides of the window triangle. Firstly it is clear that $L_{12}$ = $L_2$. Then the projected distance (top view) $d$ between $L_1$ and $L_2$ is $h/ tan\,\theta$, where $h$ is the sapphire wafer thickness. Since $\beta = 30°$, we have $L_{11} = d \times \tan\beta = \sqrt{3}h/ tan\,\theta = L_{13}$. Therefore $L_1 = L_{11} + L_{12} + L_{13} = L_2 + 2\sqrt{3}h/ tan\,\theta$.

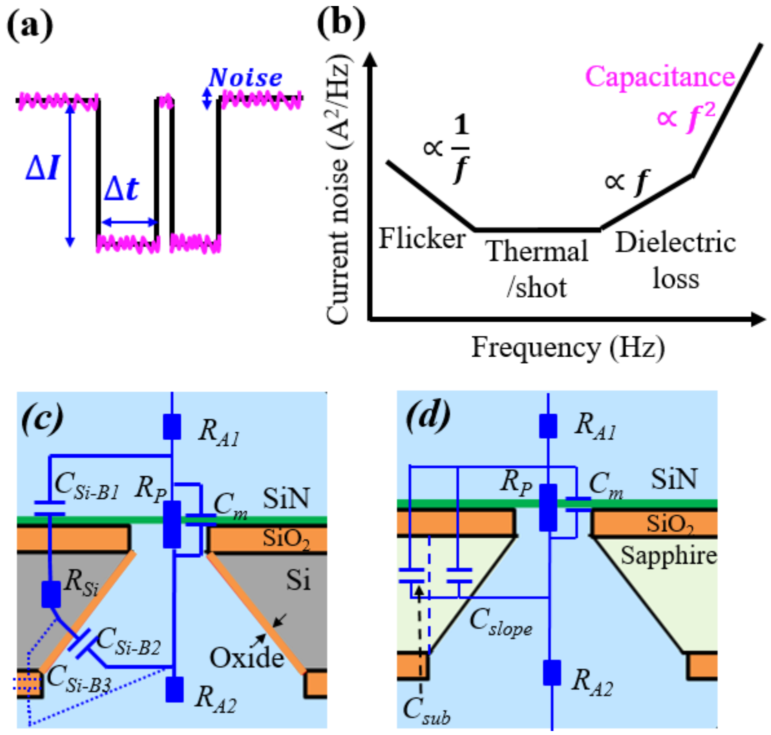


**Figure S1**. Motivation of designing low-noise solid-state nanopores in sapphire. (a) A schematic of typical DNA signals during the DNA translocating through a solid-state nanopore. $\Delta I$ is the blockade current amplitude, $\Delta t$ is the dwelling time, and the pink ripples are the current noise. (b) The current noise contribution at different frequencies. One key noise contributor at high-frequency detection is the total input capacitance. (c) The equivalent circuit of a silicon-substrate solid-state nanopore, showing the parasitic capacitance ($C_{Si\text{-}B1}$, $C_{Si\text{-}B2}$ and $C_{Si\text{-}B3}$) due to the existence of free carriers in the silicon substrate. (d) The equivalent circuit of a sapphire-substrate solid-state nanopore. No parasitic capacitance is observed due to the insulating property of the sapphire substrate. Instead, as a dielectric material, the capacitance from the thick sapphire itself ($C_{sub}$ and $C_{slope}$) are very small.

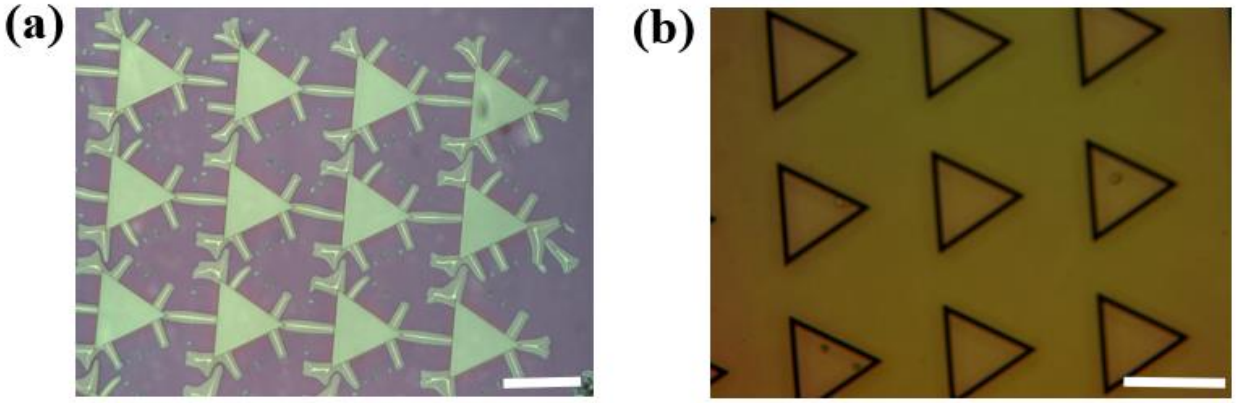


**Figure S2**. Process development to achieve crack-free $SiO_2$ mask for reliable sapphire etching. (a) An optical image of a sapphire wafer with PECVD $SiO_2$ window patterned after 1-hour sapphire etching ( hot-plate temperature set at 400 °C). Severe undercut etching was observed. Scale bar: 400 μm. (b) An optical image of the sapphire wafer with the same PECVD $SiO_2$ window after 2-hour sapphire etching (hot-plate temperature set at 450 °C), with added RCA2 cleaning and thermal annealing prior to etching. No undercut etching was observed. Scale bar: 400 μm.

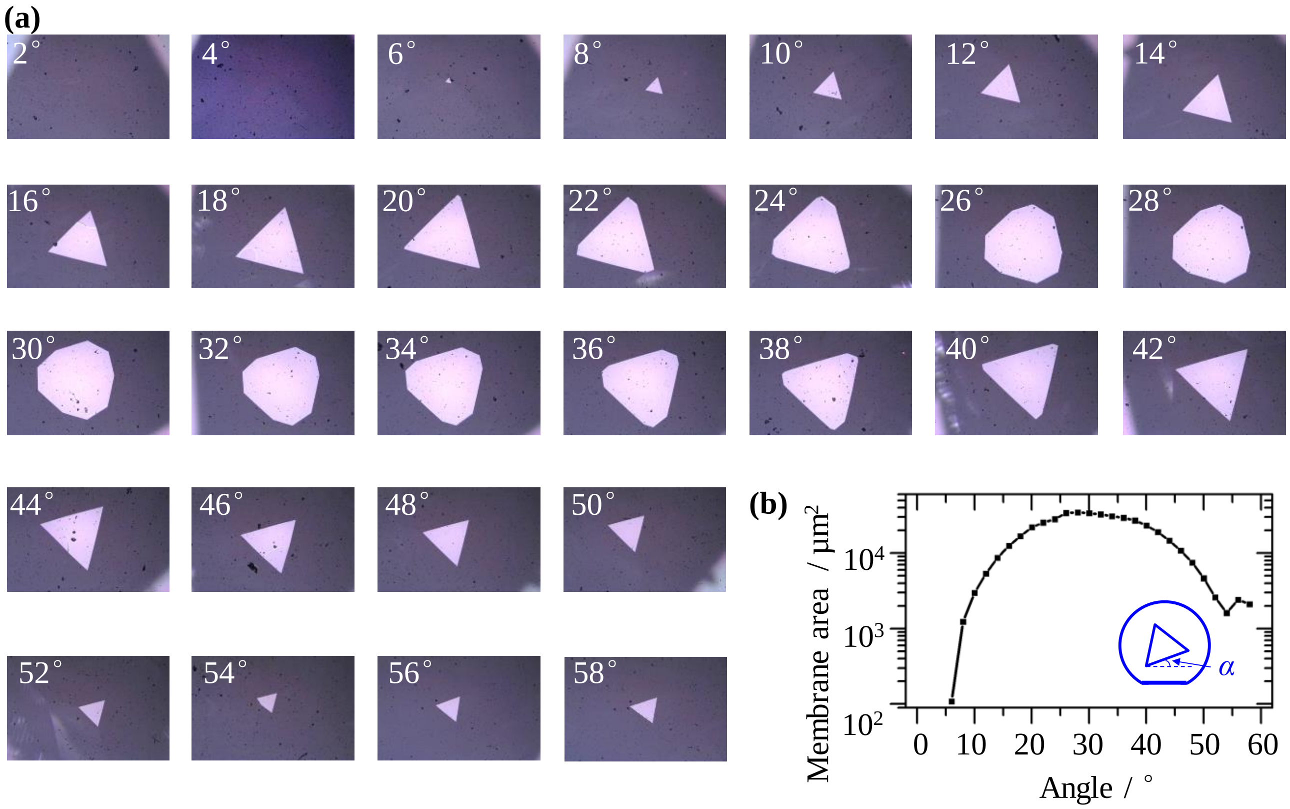


**Figure S3**. Experimental analysis of the dependence of membrane shape and size (side length $L_2$) on the alignment angle between the triangular-shaped etching windows (side length $L_1$) and the A-plane sapphire flat. (a) Optical images of the membranes. The alignment angles (*α*, indicated in figure b) between the etching window and the A-plane sapphire flat is indicated on the images. (b) The plot of the membrane area versus the alignment angle *α*. Here the etching window size length $L_2$ was fixed as about 767 μm. The sapphire was etched about 232 μm in depth (*i.e.* remaining thickness ~20 μm) .

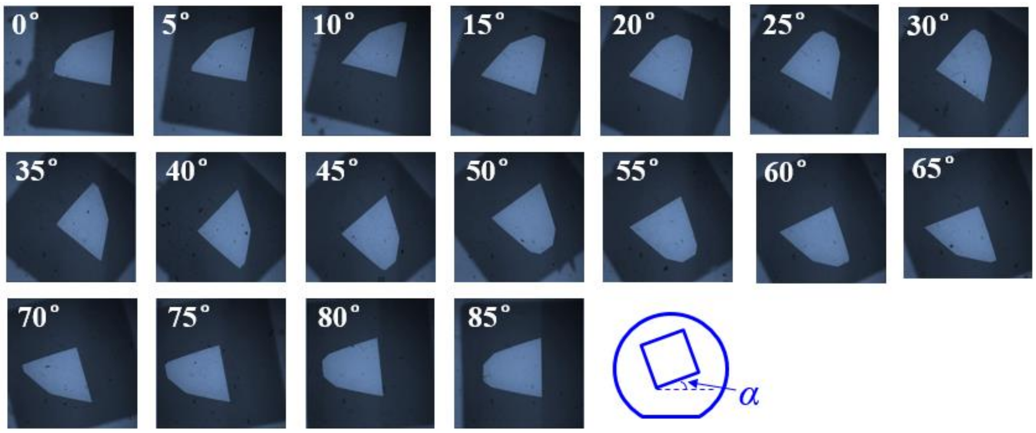


**Figure S4**. Optical images of the membranes formed on sapphire using square-shaped etching windows ($L_1$) with different alignment angles. Here the window side length $L_2$ was fixed as 800 μm. The numbers on each image indicate the alignment angles $\alpha$.

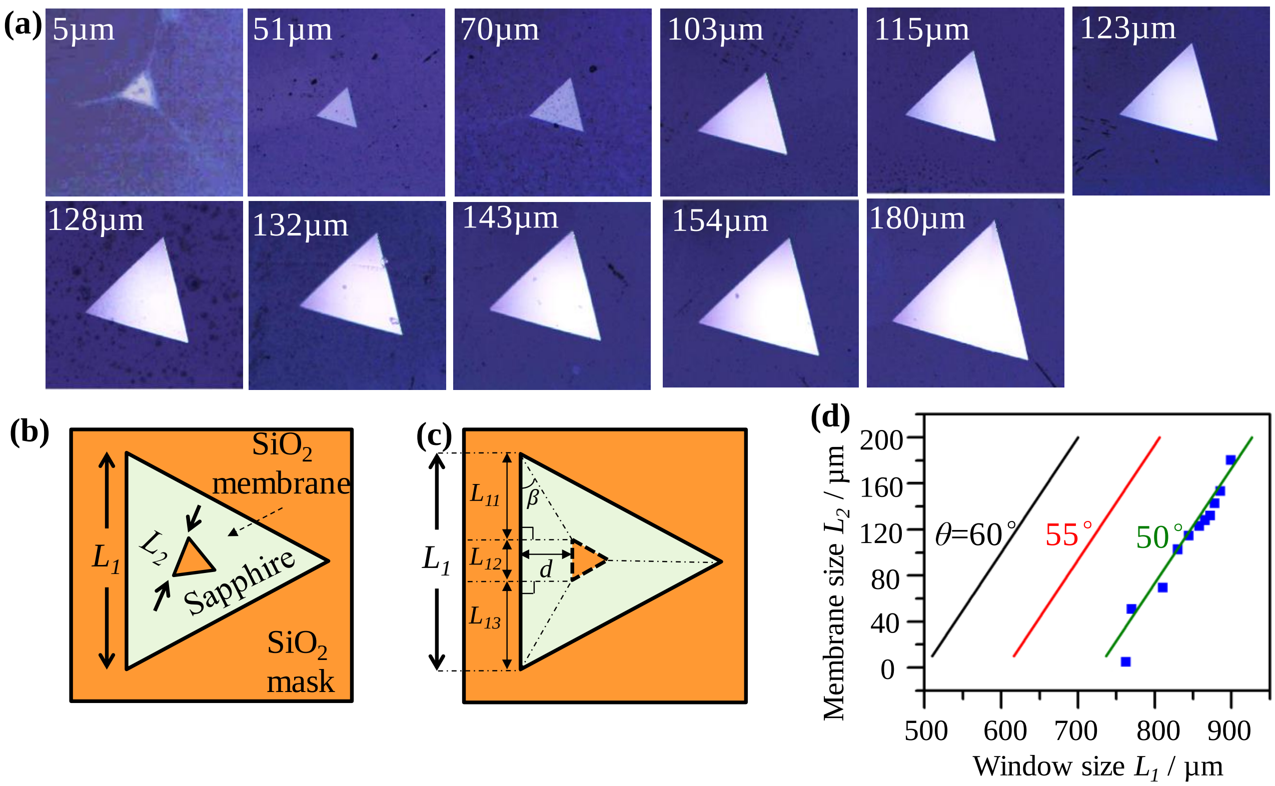


**Figure S5**. Demonstration of tuning membrane dimension by engineering the etching mask dimensions. (a) Optical images of the membranes (side length $L_1$) with different etching window sizes (side length $L_2$). The numbers on the images indicate the value of $L_2$. The magnification of the objective lens is 100x for the 5 µm triangle and 10x for others. (b) A schematic top view of a real sapphire chip after the sapphire is etched through. There is an offset angle between $L_1$ and $L_2$, attributed to facet competition during etching (c) A schematic top view showing the estimation of the effective facet angle $\theta$, by assuming the membrane sides $L_2$ parallel to mask triangle sides $L_1$ and applying the relationship $L_1 = L_2 + 2\sqrt{3}h/\tan\theta$. (d) Plot of $L_2$–$L_1$ relationship, fitted by a model assuming the sapphire etching follows an effective facet angle $\theta$, which is the angle marked in Figure 2a. The fitting indicates the effective facet angle is around 50° while $\alpha$=0°.

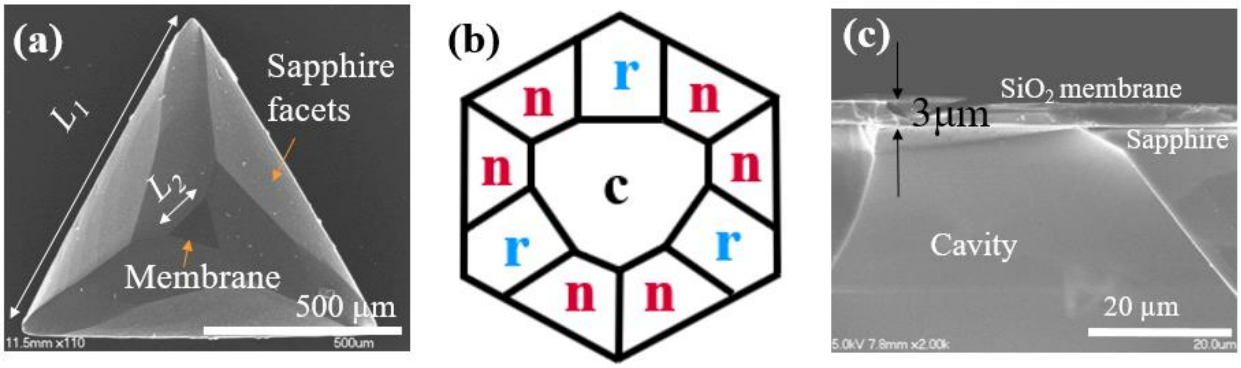


**Figure S6**. Facets of the sapphire in the cavity after sapphire etching. (a) A SEM image of the top view of the sapphire cavity. (b) The three-fold symmetric n-r-n plane system, which caused the three-fold symmetric etching facets of sapphire. (c) The SEM image of the cross section of the sapphire cavity.

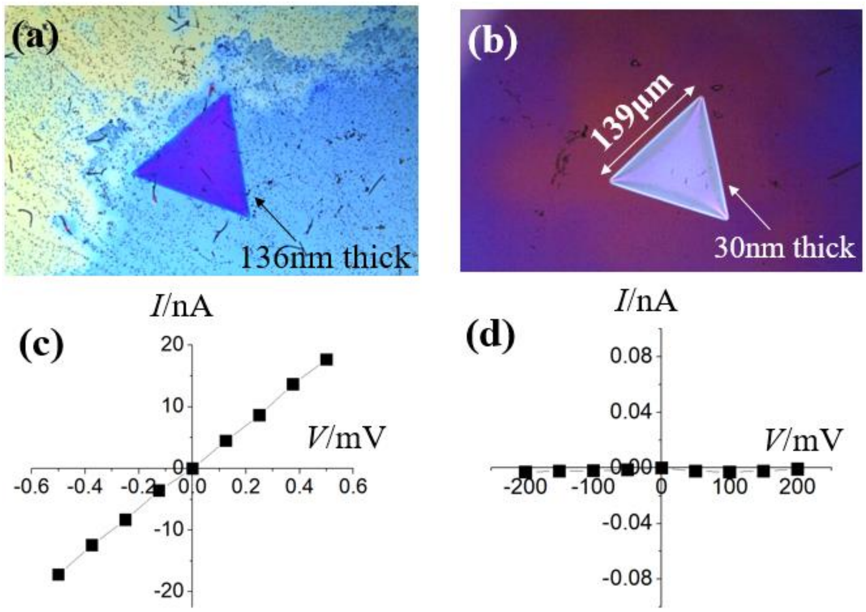


**Figure S7**. Comparison of device leakage current by different SiN thinning methods. (a) Optical image of a SiN membrane after RIE dry etching (thickness after etching: 136 nm). RIE recipe: PlasmaTherm 790 RIE Fluorine (tool), 30 W bias, 100 mTorr, $CF_4$ 50 sccm, $O_2$ 2 sccm, etching rate: 18 nm/min (b) Optical image of a SiN membrane after hot phosphoric wet etching (thickness after etching: 30 nm). (c) Current-voltage (*IV*) characteristic of the membrane in figure a in 1M KCl solution. (d) *IV* characteristic of the membrane in figure b in 1M KCl solution.

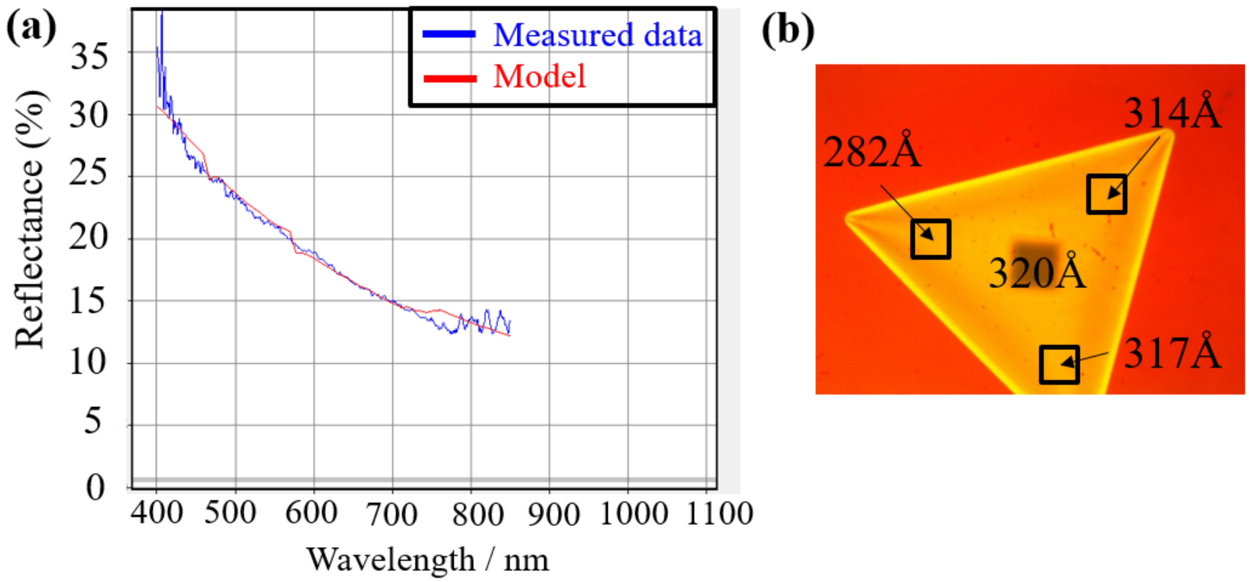


**Figure S8**. The thickness characterization of the membrane thickness using Filmetrics F40. (a) The fitting curve (red) of the measured reflectance spectrum of the SiN membrane (blue). (b) The uniformity characterization of the membrane thickness. The thickness variation of the left corner may come from the bending of the membrane, since F40 measurement is based on the reflectance of light.

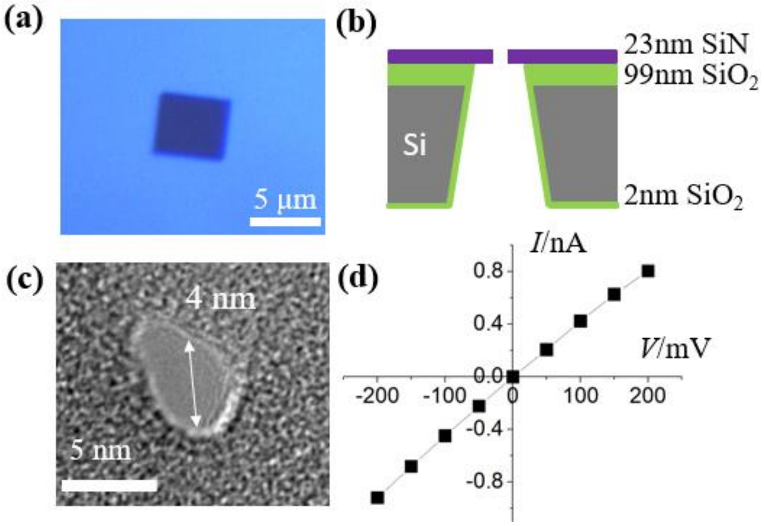


**Figure S9**. A small-membrane SiS nanopore chip for comparison. (a) The optical image of the SiN membrane (4.2 µm by 4.7 µm). (b) The schematic of the structure of this nanopore chip. (c) The nanopore drilled by TEM on the SiN membrane. (d) The *IV* curve tested by 100 mM KCl solution, showing good linearity.

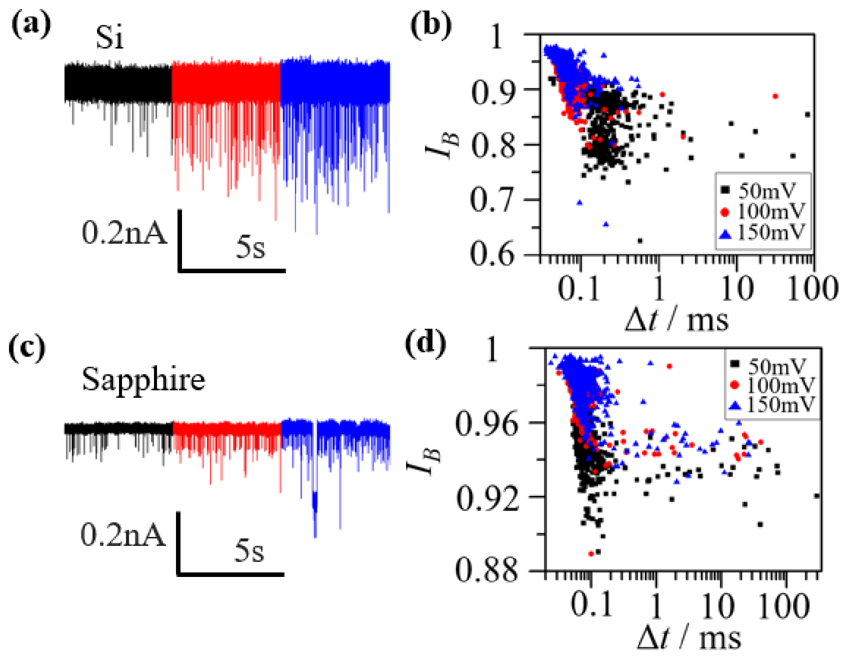


**Figure S10**. Representative 1kbp dsDNA translocation events for the SaS nanopore and the SiS nanopore under 10kHz filter bandwidth. (a) The current trace of the DNA translocation events of the SiS nanopore under different voltages (black: 50 mV, red: 100 mV, blue: 150 mV). (b) Scatter plot of the fractional blockade current $I_B$ ($=i_b/i_0$) versus the dwelling time $\Delta t$ of all the DNA events from the SiS nanopore under different voltages (black: 50 mV, red: 100 mV, blue: 150 mV). (c) The current trace of the DNA translocation events of the SaS nanopore under different voltages (black: 50 mV, red: 100 mV, blue: 150 mV). (d) Scatter plot of the fractional blockade current $I_B$ ($=i_b/i_0$) versus the dwelling time $\Delta t$ of all the DNA events from the SaS nanopore under different voltages (black: 50 mV, red: 100 mV, blue: 150 mV).

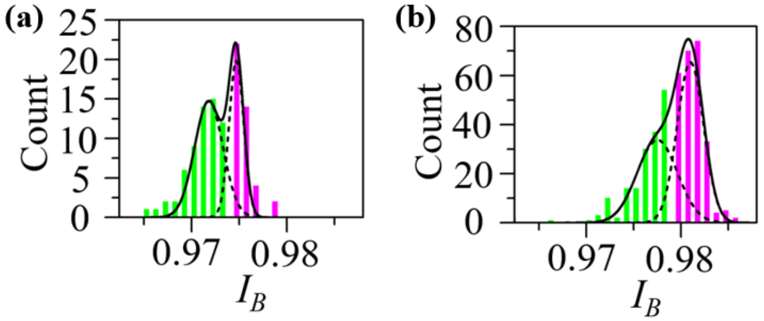


**Figure S11**. The histograms of $I_B$ from the analysis of Poly(A)$_{40}$ single-stranded (ss) DNA by the SaS nanopore under 100 mV (a) and 150 mV (b). Two distinct peaks are observed and fitted by Gaussian function, corresponding to the translocation events (green bars) and the collision events (pink bars).

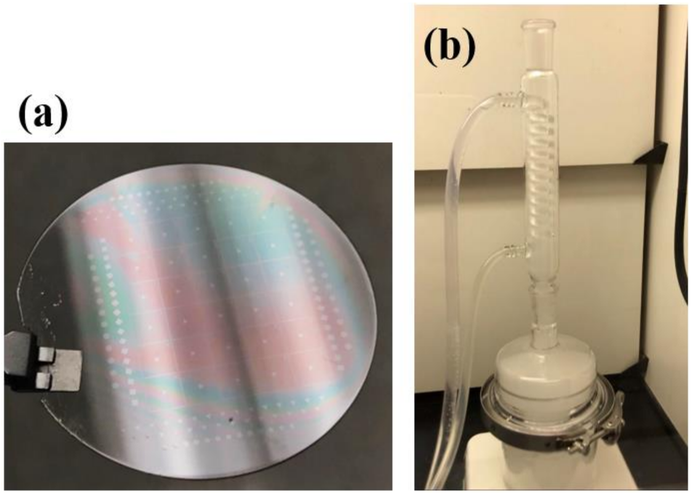


**Figure S12**. Optical graphs of: (a) A sapphire wafer with $SiO_2$ mask patterned right before the sapphire etching. (b) Our customized glassware setup used for sapphire etching: The quartz vessel and its lid are clamped together by a stainless-steel clamp. On the top, a water condenser is used to condense and recirculate the evaporated acid. The setup is hosted on a hot plate during etching. Due to safety reasons, we could not precisely monitor the solution temperatures during etching, but we used the same etching volume and monitor the hotplate temperature setting to ensure the repeatability of the etching process. We also note that commercially available etching tanks are available (https://www.imtecacculine.com/high-temperature-quartz-tank) that can automate the etching and cooling process and handle up to 25 pieces of 2-inch sapphire wafers in one batch.

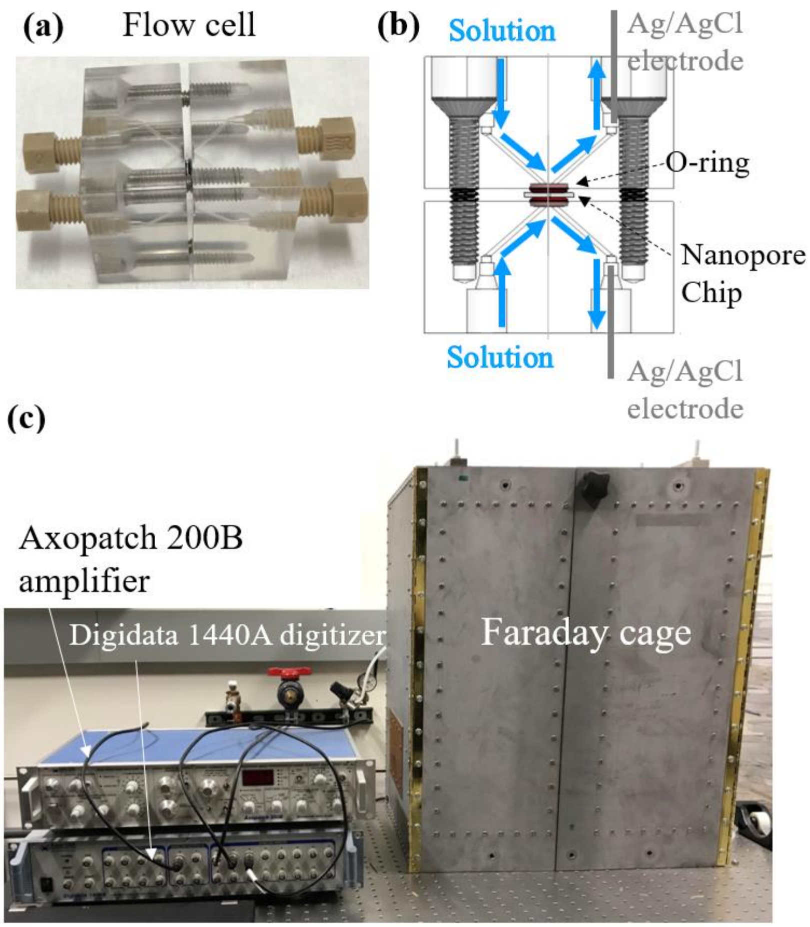


**Figure S13**. The experimental setup for the noise characterization and DNA sensing of the nanopore chip. (a) A photo of the flow cell used for providing electrolyte ambient for the nanopore chip. It is composed of two acrylic pieces drilled with fluidic channels. The two pieces are mounted together by four screws. (b) A schematic of the flow cell showing the injection of the electrolyte solution (blue path) and the mounted nanopore chip. (c) The Faraday cage used to contain the flow cell to isolate the environment noise and the Axopatch 200B amplifier with the Digidata 1440A digitizer.
