## Supporting tables for "Sapphire-Supported Nanopores for Low-Noise DNA Sensing"

**Table S1**. The calculation of the SiS nanopore chip capacitance in Figure S1c.

| Capacitance | Material | Dielectric constant ($\varepsilon$) | Area ($A$) | Thickness ($t$) | Calculation Method[b)] | Calculated capacitance |
|---|---|---|---|---|---|---|
| $C_{Si-B1}$ | SiN ($C_1$) and $SiO_2$ ($C_2$) | 6.5 ($\varepsilon_1$) and 3.9 ($\varepsilon_2$) | $(\frac{D_o}{2})^2 \times \pi - A_{m\_si}$[a)] | 23nm SiN ($t_1$) and 99nm $SiO_2$ ($t_2$) | $C_{Si-B1} = \frac{1}{1/C_1 + 1/C_2}$ <br> $C_1 = \varepsilon_1 \cdot \varepsilon_0 \cdot A/t_1$ <br> $C_2 = \varepsilon_2 \cdot \varepsilon_0 \cdot A/t_2$ | **1384pF** |
| $C_{Si-B2} + C_{Si-B3}$ | $SiO_2$ | 3.9 | $\approx (\frac{D_o}{2})^2 \times \pi$ | 2nm | $C_{Si-B2} + C_{Si-B3} = \varepsilon \cdot \varepsilon_0 \cdot \frac{A}{t}$ | **70106pF** |
| $C_m$ | SiN | 6.5 | $A_{m\_si}$ | 23nm | $C_m = \varepsilon \cdot \varepsilon_0 \cdot \frac{A}{t}$ | **0.049pF** |
| $\boldsymbol{C_{total}}$ | | | 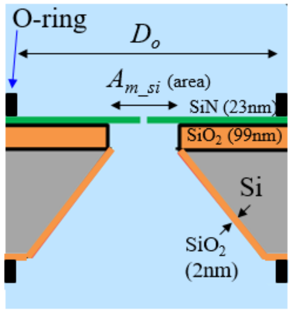 | | $C_{total} = C_m + C_{Si-B1} \| \| (C_{Si-B2} + C_{Si-B3})$ | **1360pF** |



The SiS nanopore (from SiMPore) has 4.2×4.7 μm$^2$ wide, 23 nm thick membrane.

a) $D_o$ (2400 μm) is the real o-ring opening diameter, which was measured by stamping an o-ring pattern on a piece of paper using some low-moisture ink. $A_{m\_si}$ is the membrane area, calculated by 4.2 times 4.7 μm (side width and side length, respectively).

b) $\varepsilon_0$ is the vacuum permittivity, 8.854 pF/m. The variables are shown in the schematic.

**Table S2**. Comparison of representative low-noise solid-state nanopore chips reported from literature.

| **Methods** | **Schematic** | **Substrate/ membrane** | **Membrane area/ thickness** | **Chip capacitance** | **RMS noise current** | **PSD plot (y-axis: Power ($pA^2/Hz$)** | **Membrane fabrication comments** |
|---|---|---|---|---|---|---|---|
| HF etching glass & SiN membrane transfer[1] | 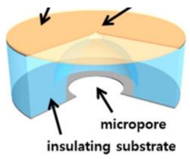 | Glass/ SiN | 25$\mu m^2$/ 20nm | 70pF (measured) | 12.58pA @10kHz, 4.5nA | 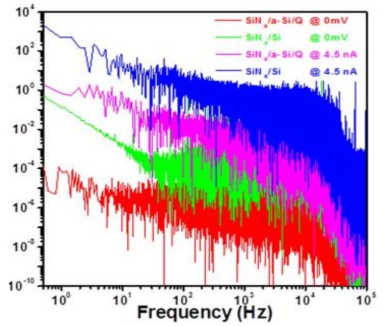 | ■ Manual transfer of SiN membrane: potentially low-yield and low-throughput. |
| HF etching glass & SiN membrane transfer[2] | 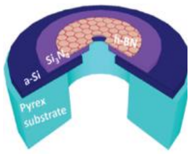 | Glass/ h-BN | 0.008$\mu m^2$/ 2-3nm | 5-10pF (measured) | 4.3pA @ 10kHz, 0nA; 12.8pA @100kHz , 0nA | 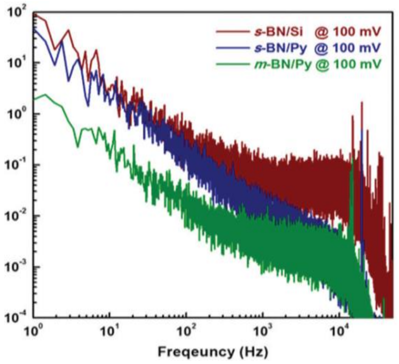 | ■ Manual transfer of SiN membrane: potentially low-yield and low-throughput.<br>■ Additional FIB drilling to make the very small h-BN window: potentially low-throughput. |
| Two-step HF etching glass & Silicone painting[3] | 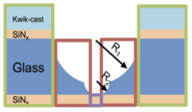 | Glass/ Graphene | 0.071$\mu m^2$/ 0.34nm | 0.69-1.65pF (measured) | NA | NA | ■ Multi-step lithography and etching: complex and potentially low-throughput.<br>■ Manual painting of silicone: potentially low-throughput and low-yield. |

**Table S2**. (Continued).

| Methods | Schematic | Substrate/ membrane | Membrane area/ thickness | Chip capacitance | RMS noise current | PSD plot (y-axis: Power ($pA^2/Hz$) | Membrane fabrication comments |
|---|---|---|---|---|---|---|---|
| Laser density modification of fused silica[4] | 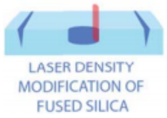 | Fused silica/ SiN | 314μm²/ 30nm | 2.2pF (measured)/ 0.75pF (calculated) | 18.7pA @10kHz | 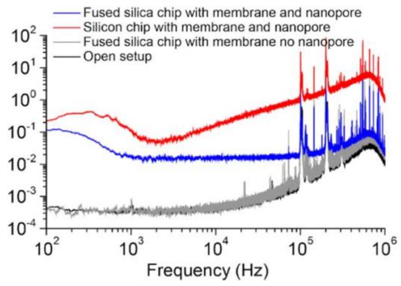 | ■ Accelerated etching of laser-exposed fused silica potentially feasible for wafer-scale fabrication. ■ Membrane size variation seems to be large. ■ Impact of laser use on throughput and cost unknown. |
| Customized amplifier & EBL & Silicone painting[5] | 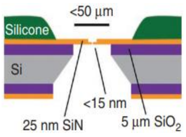 | Si/ SiN | 2500μm²/ 25nm (center: 0.25μm²/ 10-15nm) | 6pF (calculated) | 7.2pA @10kHz; 12.9pA @100kHz (Bias info: NA) | 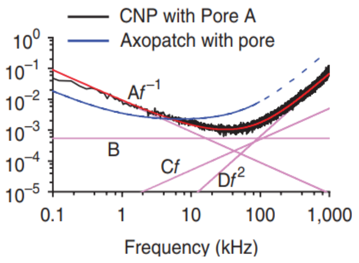 | ■ Manual painting of silicone: potentially low-throughput. ■ Extra nanofabrication for a small membrane: complex and potentially low-throughput. |

**Table S2**. (Continued).

| **Methods** | **Schematic** | **Substrate/ membrane** | **Membrane area/ thickness** | **Chip capacitance** | **RMS noise current** | **PSD plot (y-axis: Power ($pA^2/Hz$)** | **Membrane fabrication comments** |
|---|---|---|---|---|---|---|---|
| Manual painting and bonding & EBL[6] | 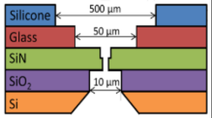  | Si/ SiN | 100-1600µm²/ 50nm (center: 0.014µm²/ sub 10nm) | 1.9-5.8pF (measured) | 119-149pA @1MHz (Bias info: NA) | 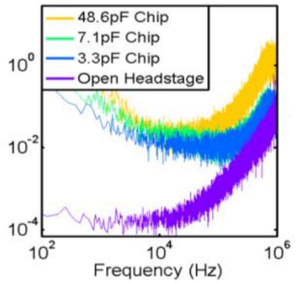  | ■ Manual painting and bonding: potentially low-throughput. ■ Extra nanofabrication for small membrane: complex and potentially low-throughput. |
| This work: Sapphire substrate - Anisotropic etching | 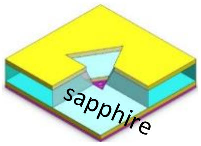  | Sapphire/ SiN | 2002µm²/ 30nm | 10pF (measured)/ 5.4pF (calculated) | 4.7pA @10kHz, 50mV<br><br>17.7pA @100 kHz, 50mV | 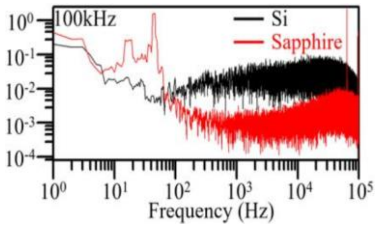  | ■ Anisotropic etching of sapphire, simple structure and fabrication scheme. ■ Batch processing compatible, potentially inexpensive and high-throughput. |

**Table S3**. The calculation of the SaS nanopore capacitance in Figure S1d.

| Capacitance | Material | Dielectric constant ($\varepsilon$) | Area ($A$) | Thickness ($t$) | Calculation Method[e)] | Calculated Capacitance |
|---|---|---|---|---|---|---|
| $C_{sub}$ | Sapphire | 9.3 | $(\frac{D_o}{2})^2 \times \pi - A_{cav\_sa}$[a)] | 250μm | $C_{sub} = \varepsilon \cdot \varepsilon_0 \cdot \frac{A}{t}$ | **1.4pF** |
| $C_{slope}$ | Sapphire | 9.3 | $L_2$ edge to $L_1$ edge[b)] | 250μm | $C_{slope} = \int_{L2\ edge}^{L1\ edge} \varepsilon \cdot \varepsilon_0 \cdot \frac{A\,(x)}{t(x)} dx$ | **0.2pF** |
| $C_{m\,(case\ 1)}$ | SiN | 6.5 | $(L_{2\,(case\ 1)}/2)^2 \times \sqrt{3}$[c)] | 30nm | $C_{m\,(case\ 1)} = \varepsilon \cdot \varepsilon_0 \cdot \frac{A}{t}$ | **3.8pF** |
| $C_{m\,(case\ 2)}$ | 2D materials (*e.g.* hBN, $MoS_2$, $WS_2$) | 3~4, 7.2, 6.5[7] | $(L_{2\,(case\ 2)}/2)^2 \times \sqrt{3}$[d)] | 2nm | $C_{m\,(case\ 2)} = \varepsilon \cdot \varepsilon_0 \cdot \frac{A}{t}$ | **0.2~0.3pF** |
| $\boldsymbol{C_{total}}$ | | | 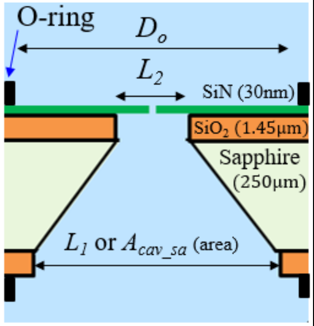 | | $C_{total} = C_{sub} + C_{slope} + C_m$ | **5.4pF (case 1)**<br>**1.8~1.9pF (case 2)** |

$C_{m\ (case\ 1)}$: experimental values. $C_{m\ (case\ 2)}$: hypothetical future integration with 2D materials.

[a] $D_o$ (2400 µm) is the o-ring opening diameter, and $A_{cav\_sa}$ is the projected area (top view) of the backside cavity of the chip, calculated as $(L_1/2)^2 \times \sqrt{3}$ ($L_1$ = 762 µm).

[b] Projected area (top view) between $L_2$ edge to $L_1$ edge.

[c] $L_{2\ (case\ 1)}$ = 68 µm, [d] $L_{2\ (case\ 2)}$ = 5 µm.

[e] $\varepsilon_0$ is the vacuum permittivity, 8.854 pF/m. The variables are defined in the schematic.
